# Developmental transcriptomes predict adult social behaviors in the socially flexible sweat bee, *Lasioglossum baleicum*

**DOI:** 10.1101/2023.08.14.553238

**Authors:** Kennedy S. Omufwoko, Adam L. Cronin, Thi Thu Ha Nguyen, Andrew E. Webb, Ian M. Traniello, Sarah D. Kocher

## Abstract

Natural variation can provide important insights into the genetic and environmental factors that shape social behavior and its evolution. The sweat bee, *Lasioglossum baleicum*, is a socially flexible bee capable of producing both solitary and eusocial nests. We demonstrate that within a single nesting aggregation, soil temperatures are a strong predictor of the social structure of nests. Sites with warmer temperatures in the spring have a higher frequency of social nests than cooler sites, perhaps because warmer temperatures provide a longer reproductive window for those nests. To identify the molecular correlates of this behavioral variation, we generated a *de novo* genome assembly for *L. baleicum*, and we used transcriptomic profiling to compare adults and developing offspring from eusocial and solitary nests. We find that adult, reproductive females have similar expression profiles regardless of social structure in the nest, but that there are strong differences between reproductive females and workers from social nests. We also find substantial differences in the transcriptomic profiles of stage-matched pupae from warmer, social-biased sites compared to cooler, solitary-biased sites. These transcriptional differences are strongly predictive of adult reproductive state, suggesting that the developmental environment may set the stage for adult behaviors in *L. baleicum*. Together, our results help to characterize the molecular mechanisms shaping variation in social behavior and highlight a potential role of environmental tuning during development as a factor shaping adult behavior and physiology in this socially flexible bee.

## Introduction

Systems that encompass natural variation in behavior can help to unmask the genetic and environmental factors that shape these complex traits. The halictine bee, *Lasioglossum baleicum*, is a socially flexible bee species capable of producing both eusocial and solitary nests (Figure 1). This sweat bee is common in the temperate regions of Japan, where nests at lower elevations and latitudes are primarily eusocial while high elevation populations produce all solitary nests (Hirata & Higashi, 2008). Beyond this clinal variation in eusociality, some populations in Northern Japan include a mix of eusocial and solitary nests at the same site, with social nests found primarily in sunny patches of the nesting aggregation and solitary nests found primarily in the shade (Cronin & Hirata, 2003; Hirata & Higashi, 2008; Yagi & Hasegawa, 2011). Thus, *L. baleicum* females can produce both eusocial and solitary nests, and this variation can occur both across different populations or within a single population.

**Figure 1.**
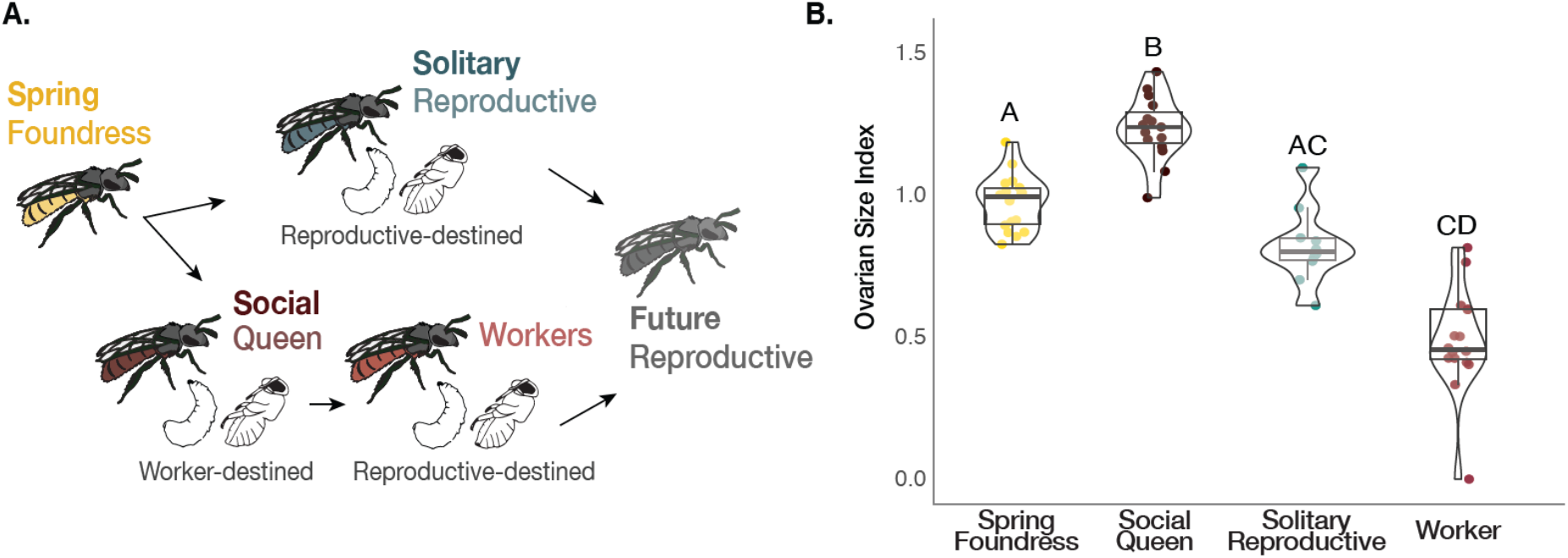
Lasioglossum baleicum is a socially flexible sweat bee. A. In the spring, individual females (spring foundresses) can either reproduce solitarily or form eusocial nests with workers in the summer (Cronin & Hirata, 2003). This variation is dependent on environmental conditions – females tend to produce eusocial nests in warmer patches and solitary nests in cooler patches of an aggregation near Sapporo in Hokkaido, Japan. B. Comparisons of the ovarian size indices (length of the longest oocyte [mm] / intertegular distance [mm]) reveal that social queens (n=15) have the greatest ovarian activation (Kruskal-Wallis*: X_3_^2^* = 51.78, *P* = 3.34e-11). Solitary reproductives (n=11) are intermediate, and workers (n=17) had the least developed ovaries. Letters represent significant differences according to Dunn’s post-hoc tests.

The behavioral variation encompassed by *L. baleicum* is likely to be adaptive. Eusocial nests have almost a 9-fold increase in reproductive success compared to solitary nests, and the inclusive fitness benefits to workers have been calculated to be higher than those of their solitary counterparts (Yagi & Hasegawa, 2012). Despite these reproductive benefits, eusociality may not always be the most successful strategy. Solitary nests are more common than eusocial ones both at higher elevations and at cooler, shady sites (Hirata & Higashi, 2008). The cooler temperatures in these regions are likely to result in shorter seasonal windows where temperatures are warm enough for bees to forage and for brood to develop but limit reproduction to one brood per season (Kocher et al., 2014). While solitary females produce the next generation of reproductives, the eusocial strategy requires an initial investment in worker production followed by the production of a second, reproductive brood. Thus, in cooler areas, the developmental windows may not be sufficiently long for eusocial nests to complete their reproductive cycles (Kocher et al., 2014). As a result, there appears to be a trade-off between the fitness benefits associated with a eusocial reproductive strategy that initially invests in the production of workers versus the amount of time required for a nest to produce the next generation of reproductive females. This trade-off and the importance of season length as a driver of social evolution has been described and documented more broadly across many insects (Cronin & Schwarz, 1999; Davison & Field, 2016, 2018; Field, Paxton, Soro, Craze, & Bridge, 2012; Kocher et al., 2014; Plateaux-Quénu, Plateaux, & Packer, 2000; Schwarz, Richards, & Danforth, 2007; Sheehan et al., 2015).

Whether an adult female remains in a nest as a worker or disperses to become a solitary reproductive is largely flexible in adulthood (Breed et al., 1978; Brothers & Michener, 1974; Steitz & Ayasse, 2020). In *L. baleicum* and all social halictines and paper wasps, females are capable of reproduction throughout their lives (i.e., they are totipotent; Hirata & Higashi, 2008). In the absence of a queen, workers can dynamically activate their ovaries and take over as a new dominant, reproductive female in the nest. As a result, most work in sweat bees has focused on the behavioral interactions and chemical signals experienced by adults within the nest. However, developmental factors are also likely to influence adult behavior and physiology, including differences in nutrition (Kapheim et al., 2011; Lawson et al., 2017) and temperature (Dehaan et al., 2022; Hirata & Higashi, 2008). Reproductive females typically eclose with greater fat stores (Brand & Chapuisat, 2012) and, in many lineages, these females are often larger than workers. Thus, the social flexibility observed in *L. baleicum* is likely to result from a complex interplay of developmental and social environments.

To better understand the factors that shape and potentially maintain social flexibility in *L. baleicum*, we characterized the environmental and molecular correlates of the different behavioral strategies found within a single population of this species located near Sapporo, Japan. In this population, females either produce a solitary nest with a single, mixed-sex brood or eusocial nests with an initial female-biased worker brood followed by a mixed-sex reproductive brood, typically separated by an inactive period of several weeks (Figure 1a; Cronin & Hirata, 2003; Davison & Field, 2018). We first examined the relationship between soil temperature and nest social structure throughout the active season. Then, we characterized differences in gene expression patterns across key developmental timepoints and social phenotypes. To this end, our study highlights the interplay between environmental variation and individual physiology in modulating social behavior in this unique, socially flexible bee.

## Materials and Methods

### Study organism and sample collections

*L. baleicum* individuals were collected from Nishioka park in Japan (near Sapporo, western Hokkaido; alt. 150 m, 141°35′ E, 43°00′ N). Temperature probes (IBS-TH2, INKBIRD) were placed in the soil at approximately 20cm depth to approximate the position of the brood clusters. Data were collected during two trips: the first in June 2021 and the second in July 2021.

Nests in this aggregation contain a mix of solitary and eusocial nesting strategies (Figure 1a). Solitary nests consist of a single, reproductive female solely responsible for provisioning her brood. This brood is a mix of reproductive males and females produced in a single, continuous bout throughout the season. Eusocial nests consist of multiple females, with a single queen and an average of 2-3 workers provisioning the brood. Social queens produce two broods: first a female-biased brood of workers followed by a mix of reproductive males and females (gynes). Like many other halictine bees, males disperse after eclosion. All females are morphologically similar, and eusocial workers are capable of mating and reproduction in the absence of a queen. Many mid-season nests contain multiple adult females, with some evidence of nest swapping among workers (Yagi & Hasegawa, 2012).

Samples were collected at two different timepoints. The first collection (spring) was done on June 16, 2021, when nests contained spring foundresses, pupae from brood one, and no adult offspring. The second collection (summer) was July 28, 2021, when nests contained founding females (solitary reproductives or queens), adult workers (social nests only) and developing brood (eggs, larvae, pupae). Collections began at 04:00 by blocking the nest entrance with a plastic cup to ensure none of the adults escaped when foraging time approached. During excavation, white talcum powder was injected into each individual nest entrance with a Pasteur pipette to distinguish individual nest tunnels and architecture, facilitating collection of individuals belonging to the focal nest. To assess environmental correlates, temperature probes were placed in the soil at the field site on collection days to record soil temperatures for 72 hours. Excavated nests were proximal to these temperature probes and this information was recorded in the sample metadata. Individual bees were placed in labeled 1.5ml cryoresistant microcentrifuge tubes and immediately flash-frozen in liquid nitrogen.

We collected a total of 62 adults and 22 pupae for this study from 34 nests: eusocial queens (n = 15), spring foundresses (n = 19), solitary reproductives (n = 11), eusocial workers (n =17), spring worker-biased female pupae (n = 11), and spring solitary-biased female pupae (n = 11; Table S1). Pupae used for gene expression analyses were stage-matched based on cuticle and eye pigmentation; we selected late-stage pupae with no pigmentation on cuticles and light purple eyes (Tian & Hines, 2018). We did not include males in this study as they do not contribute to nest activities and disperse soon after they emerge.

### Ovary dissections, phenotypic measurements, and caste assignments

Abdomens were dissected in RNALater ICE under a Leica M125 microscope, and pictures of ovaries were captured with the Leica MC190 HD camera. We measured intertegular distance (IT span) as a proxy for body size and measured the length of the longest oocyte to quantify ovarian development. These measures were taken for all adult bees used for gene expression analysis in order to infer caste differences among the adults (as in Kapheim et al., 2012). Data are included in Table S1.

We next calculated the ovarian index for each bee by taking the measurement of the longest oocyte and dividing it by IT span (Saleh & Ramírez, 2019). All individuals were measured blindly to prevent any bias in body size measurements. Ovary length was quantified using ImageJ v1.53n. Because ovary length distributions deviated from normality, we implemented Kruskal-Wallis tests followed by post-hoc Dunn’s pairwise tests (the adjusted p-values adjusted for multiple correction factors) to assess relationships between body and ovary size differences across behavioral groups. All analyses were carried out in R (R Core Team, version 4.1.2).

### Genome sequencing and assembly

A *de novo* genome for *L. baleicum* was generated using 10x genomics linked-reads technology. DNA was extracted from a single, frozen individual collected from Nishioka Park using a Qiagen Genomic Tip kit (Qiagen, USA, Catalog #10223) to isolate high molecular weight DNA. Libraries were sequenced to approximately 40x coverage. An assembly with a scaffold N50 of 1.9Mb was generated using Supernova v2.1.1 (Weisenfeld et al., 2017) and subsequent filtering and other quality control methods followed those of (Jones *et al* 2023). The assembled genome is available on GenBank, accession GCA_022376115.1. Genome assembly statistics and completeness are presented in Table S2 (compleasm/miniBUSCO;Huang & Li, 2023).

### Gene annotations

Gene annotations were initially predicted by BRAKER v2.1.6 (Brůna et al., 2021) using a two-run procedure. The first run derived predictions from protein hints using the *L. baleicum* repeat-softmasked assembly. The protein hints were generated by mapping a collection of orthologous arthropoda proteins downloaded from OrthoDB (Kriventseva et al., 2019) to the *L. baleicum* assembly using ProtHint v2.6.0 (Tom et al., 2020). The second run derived predictions from mapped RNAseq reads to the *Lasioglossum baleicum* repeat-softmasked assembly. RNAseq reads were aligned using STAR v2.7.5c (Dobin et al., 2013). The results of the two-run predictions were then combined using TSEBRA v1.0.2 (Gabriel et al., 2021) and fed into MAKER v3.01.04 (Cantarel et al., 2008) as ab-initio predictions. Additional evidence included protein sequences from *Bombus impatiens* (GCF_000188095.3_BIMP_2.2, UP000515180) and *Apis mellifera* (Amel_HAv3.1, UP000005203) to give evidence of protein homology and reconstructed transcripts generated by PASA v2.5.0 (Haas et al., 2003) from RNAseq reads assembled by Trinity v2.13.2 (de novo assembly and genome-guided; Grabherr et al., 2011) to provide EST evidence. We also included the always_complete and correct_est_fusion options. The output of MAKER includes both predictions that overlap provided evidence (e.g., protein homology, EST evidence, etc.) and non-overlapping predictions. To confirm if non-overlapping predictions should be included in our annotations, we used InterProScan v5.52-86.0 (Jones et al., 2014) to identify those predictions with evidence of InterPro families. Finally, we reran MAKER with both the evidence-overlapping and InterPro-overlapping predictions, the resulting annotations were then used in this analysis. BUSCO analyses (Simão et al., 2015) were then conducted to assess the completeness of gene annotations (Table S2).

### Tissue dissections and RNA extractions

Heads were immersed in a dry ice/ethanol bath, and the frons and mandibular regions were removed to expose the brain. Then, tissues were submerged in either 200uL (heads) or 400uL (abdomens) of RNAlater ICE (Invitrogen AM7030) and stored overnight at −20^°^C. The following day, tissues were dissected in RNAlater ICE and stored at −80^°^C. For RNA extraction, tissues were homogenized with a Qiagen TissueLyser, and mRNA was extracted using magnetic mRNA isolation with Dynabeads (Invitrogen, 61011) according to the manufacturer’s protocol. Cleaned mRNA was quantified with a Qubit high sensitivity (HS) assay kit (Invitrogen, Q32852) and the integrity of RNA was estimated by electrophoresis using TapeStation 2200 with High Sensitivity D5000 ScreenTape (Agilent Technologies).

### cDNA library preparation and sequencing

Individual libraries were constructed using NEBNext Ultra II Directional Prep Kit for Illumina (NEB #7760, New England Biosystems, Ipswich, Massachusetts) and sequenced at the Princeton Genomics Core on an Illumina NovaSeq 6000 with v1.5 reagent kits with paired, 150bp reads and approximately 10 million reads per library. Libraries were pooled, sequenced on two flowcells, and then reads were combined for downstream analyses. RNA extractions, library preparations, and sequencing were randomized with respect to tissues and sex to avoid batch effects. One sample was excluded from downstream analyses due to library preparation failure. Raw transcriptomic reads are available from NCBI SRA archive under bioproject ID PRJNA884745.

### Quality filtering, read mapping and quantification

A MiSeq run was conducted for initial library quantification and quality controls. Libraries were then re-pooled and rebalanced before NovaSeq sequencing. The resulting fastq files were demultiplexed using deML, a maximum likelihood demultiplexing algorithm that allows probabilistic sample assignments, with default parameters and accounting for one set of reverse-complement indices (Renaud et al., 2015). Filtering was performed to remove any remaining adapter sequences using fastp (Chen et al., 2018). Illumina strand-specific paired-end sequences from the NovaSeq reads were quality-filtered using fastp with a 4-bp window size, and a quality threshold of 20 with an option to detect and remove adapter for paired end sequences. All reads below 50 bp were trimmed. Read qualities were calculated with FastQC (bioinformatics.babraham.ac.uk/486 projects/fastqc/). We aligned the filtered FASTQ sequences to their respective reference genome, *Lasioglossum baleicum*, with STAR v2.7.5c (Dobin et al., 2013). Subsequent fragment counts for our annotated genes were then generated using featureCounts v2.0.1 (Liao et al., 2014). Gene expression levels were quantified using exon annotations and summarized at the gene level (Table S3).

### Differential gene expression analysis

#### Global expression patterns and normalization

Analyses of differential gene expressionamong behavioral groups was carried out with the DESeq2 package in R (Love et al., 2014). To account for hidden batch effects, we used surrogate variable analysis (SVA) to identify and estimate the unknown variations (Leek & Storey, 2007). Samples with low coverage (<2 million reads) were excluded from downstream analyses, and five brain tissue and eight fat body tissues were filtered out using these criteria. All the brain tissues of spring pupae passed the threshold except one pupal fat body tissue, which was subsequently excluded. To avoid problems of estimating dispersion due to low expressed features we filtered out rows that had normalized gene counts of less than 2 in at least 10 samples. After filtering, there were 11854 adult brain genes, 10665 adult fat body genes, 11248 pupal brain genes, and 10342 pupal fat body genes that remained for analysis. Our final dataset included 54 adult brain samples, 55 adult fat body samples, 22 pupal brain samples and 19 pupal fat body samples that passed the filter and were used for downstream analysis. Differentially expressed genes (DEGs) were called in DESeq2 using a contrast approach to test for behavioral effect on gene expression in all the genes that passed the filtering threshold in both brain and fat bodies. To correct for multiple testing, we employed a Benjamini-Hochberg adjustment (Benjamini & Hochberg, 1995). Genes with a false discovery rate (FDR) < 0.001 were considered as differentially expressed.

A principal components analysis was used to visualize global patterns in gene expression among samples and tissues. Expression heat maps were constructed using the “heatmaps” package in R (Perry, 2022). Overlaps of DEGs were inspected using intersection plots that were generated by UpsetR package 1.4.0 (Conway et al., 2017). Volcano plots comparing the fold expression to the adjusted p-values to support the differential expressions were produced with the “Enhanced volcano” R package (Kevin blighe, 2022). Normalized counts were used to produce hierarchical heatmaps for brain and fat body tissues using the “heatmap’’ function in R.

#### Gene ontology enrichment analyses

To identify Gene Ontology (GO) terms overrepresented withing our DEGs, we first used Trininotate (Bryant et al., 2017) to assign GO terms to *L. baleicum* genes. We then performed GO analysis on DEGs from each behavioral or developmental contrast in the brain and in the fat body (separately). We used the ‘weight01’ algorithm with Fisher’s exact test in R package, TopGO (Alexa & Rahnenführer, 2007). We analyzed the top 20 terms in the Biological Process class, separating DEGs into upregulated and downregulated sets from each contrast: workers versus reproductive adult brains and the brains of worker-biased pupae from full sun sites versus reproductive-biased pupae from the shaded sites (see below for further details). A full set of contrasts and their GO enrichment results for all social forms and tissues can be found in Table S6.

### Pupal brain DEG predictions of adult behavioral state

Using normalized, variance-stabilized transformed counts of genes differentially expressed between reproductive- and worker-biased pupal brains (“pupal DEGs”), we evaluated a random forest algorithm’s classification accuracy of adult behavioral state when trained on either all 1121 pupal DEGs or random subsets of 10, 50, 100, or 500 pupal DEGs. Each classification for pupal DEGs was compared to a random, equivalently sized subset of non-pupal DEGs (“Random genes”) or pupal DEGs with adult behavioral state label shuffled (“Shuffled labels”). Each analysis was iterated 1000 times using the randomForest package in R (Liaw & Wiener, 2002).

## Results

### Body size does not vary among behavioral forms, but social queens have the largest ovaries

Intertegular distance, a proxy for body size, did not vary significantly among the queens, workers, foundresses, or solitary reproductive females (Figure S4; Kruskal-Wallis*: X_3_^2^* = 3.47, *p* =0.32). However, there were significant differences in ovarian development (measured as the ovarian index; the length of the longest oocyte/intertegular distance). Queens had the highest ovarian index compared to other groups, with solitary reproductives intermediate, and workers with the smallest index (Figure 1b; Kruskal-Wallis*: X_3_^2^*=51.78, *p=*3.34e-11). A complementary analysis considering solely the longest oocyte (not normalized to body size) produced similar results (Kruskal-Wallis*: X_3_^2^* = 52.52, *p =* 2.32e-11).

### Nest social structure is strongly correlated with spring soil temperatures

*L. baleicum* nests in large aggregations with hundreds to thousands of nests present in a single aggregation. In these aggregations, the peripheral nesting sites are shadier, cooler, and more humid (Figure 2a), and nests in these locations tend to be solitary (containing a single reproductive female). Nests constructed in the warmer, sunnier regions of the aggregation tend to be eusocial (e.g., contain reproductive queens and non-reproductive workers). Soil probes placed at each of the sites revealed that spring soil temperatures are tightly correlated with nest social structure (Figure 2b; linear regression, R^2^=0.963, p=5.08e-4). However, summer soil temperatures did not show a significant correlation with social structure of nests (linear regression, R^2^=0.429, p=0.158). We henceforth refer to the warmer spring soil temperature sites (sites 1-3) as “social-biased” sites, and the cooler spring temperature sites (sites 4-6) as “solitary-biased” sites.

**Figure 2.**
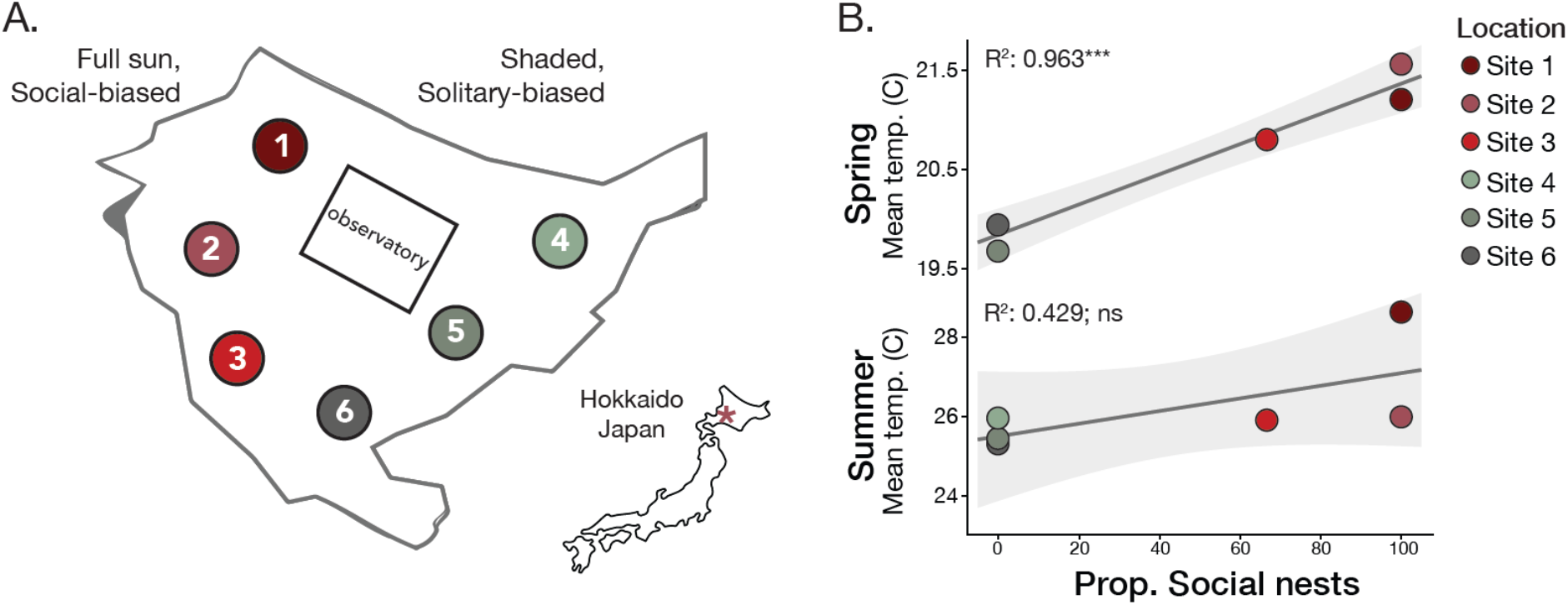
Spring soil temperatures predict nesting behavior. A) Nests were studied at six locations throughout an aggregation in Hokkaido (n=19, 3, 4, 7, 9, 6 nests excavated at each site, respectively). The aggregation is found in an open lawn surrounded by trees. Sites 1-3 receive full sun and sites 4-6 experience higher shade. B) Temperature probes placed in the soil during spring collections were strongly correlated with nesting behavior (shown here as the proportion of social nests at a given site) in the spring (linear regression, R^2^=0.963, p=5.08e-4) but not in the summer (linear regression, R^2^=0.429, p=0.158). Lines indicate the linear fit of the regressions and shading indicates the 95% confidence intervals for the model.

### Adult gene expression patterns distinguish reproductive state

We next compared the brain and fat body gene expression profiles of several behavioral phenotypes of adult females: spring foundresses, solitary reproductives, and queens and workers from social nests (Table S5; Figures. S6-S8). A principal component analysis using all transcripts that passed our initial filters separates non-reproductive workers from solitary reproductives, queens, and foundresses. Thus, the largest contribution to variation in gene expression appears to be reproductive state (Figure 3a). Similar patterns are also observed in the fat bodies (Figure S9).

**Figure 3.**
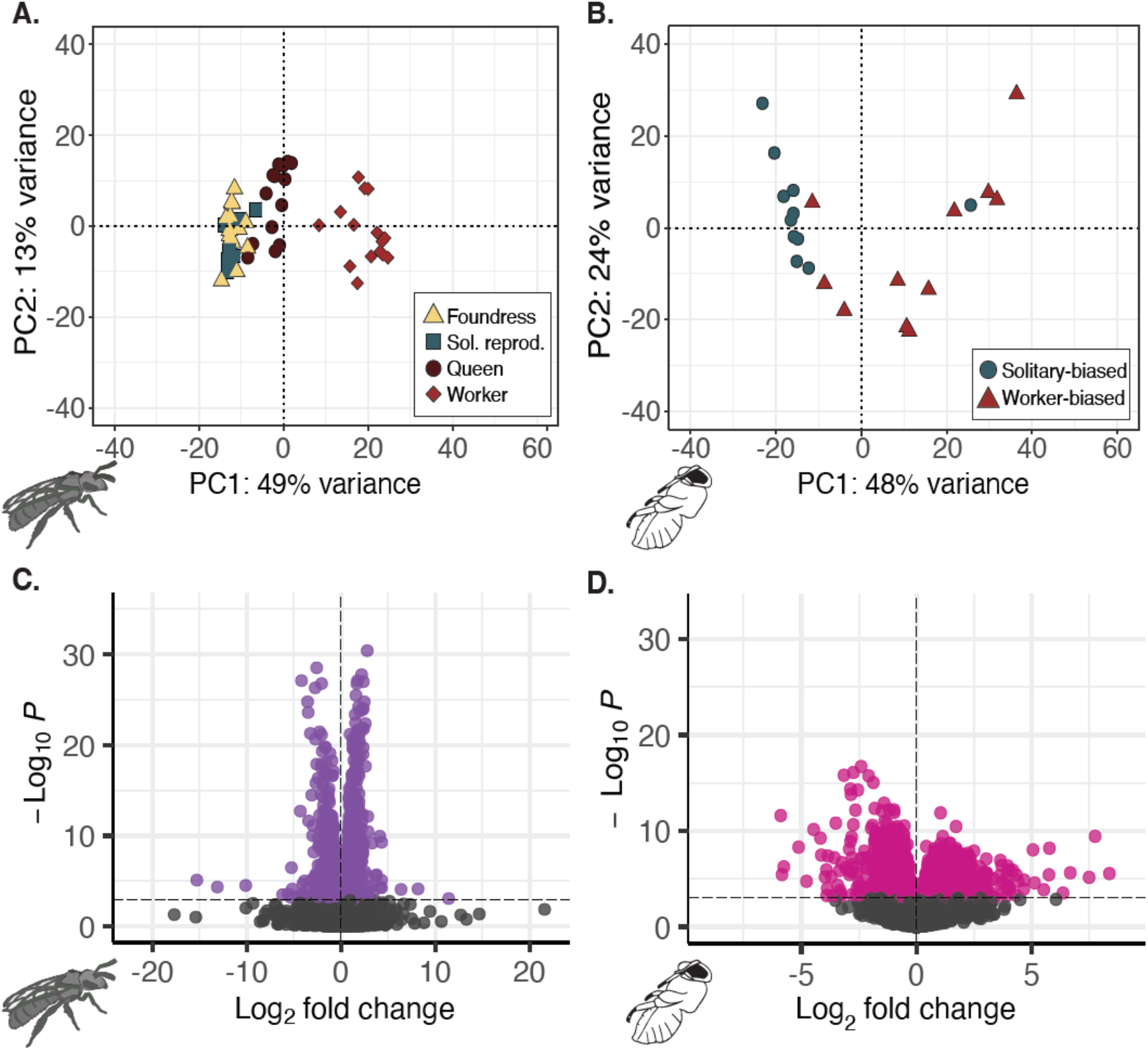
Adult and pupal brain transcriptomes reflect environmental conditions and social behavior. (A) Adult workers (n=17) have transcriptomic profiles highly distinguishable from reproductively active foundress (n=19), queen (n=15) and solitary reproductive (n=11) females. PC1 shows a strong positive correlation with ovarian development (F*_1_*_,52_=32.55, R^2^=0.37, p=5.592e-07) (B) PCA on all pupal brain transcripts discriminates between late spring pupae excavated from the social-biased (n=11) and the solitary-biased (n=11) sites. Pupae from social-biased sites are most likely worker-destined, while pupae excavated from solitary-biased sites are predicted to become reproductive females. C-D. Volcano plots highlighting DEGs of: (C) workers vs. reproductive adults (queens and solitary reproductives), and (D) Solitary-biased vs social-biased pupae (n=1121 DEGs). DEGs were determined with DEseq2 (FDR<0.001).

To gain a better understanding of which genes distinguish workers from reproductive females, we next tested for genes differentially expressed between solitary reproductives and queens versus workers and identified 1,585 differentially expressed genes in the brain (DEGs; FDR<0.001; Figure 3b). GO enrichment analyses (Table S10) reveal several Biological Processes that are associated with gene expression in workers, including protein transport and long-term memory (Figure S11a). Similarly, genes upregulated in the brains of reproductive females are enriched for transcription and translation (Figure S11b). In the fat body, genes associated with transcription and translation are upregulated in the fat bodies of reproductives (Figure S12a) while genes associated with mitochondrial function and metabolism are upregulated in worker fat bodies (Figure S12b).

We chose not to include foundresses in these analyses because they were sampled at an earlier timepoint in the season, but we note that there were 482 brain DEGs and 294 fat body DEGs between all sets of reproductive females compared to workers (Figures S7-S8). The results of all pairwise contrasts are summarized in Table S5.

### Pupal gene expression patterns are correlated with spring soil temperatures and predict adult behavior

Because spring soil temperatures were strongly correlated with the social structure of the nests (Figure 2), we were interested in comparing the gene expression profiles of stage-matched developing pupae in the shady, solitary-biased sites versus the sunny, social-biased sites. We used female pupae collected from the first spring brood. Thus, pupae located in nests from solitary-biased sites are likely to develop into solitary reproductive females while pupae collected from social-biased sites should be more likely to develop into workers.

We sequenced the brain and fat body transcriptomes, and used all of transcripts that passed our initial filters in a principal component analysis. The analysis separates pupal transcriptomes into two primary clusters along the first principal component (PC1). PC1 explains 48% of the variance among samples and is strongly correlated with nesting locality (Figure 3b; Social-biased vs Solitary-biased, *F_1_*_,20_=16.79, R^2^=0.4292, p=0.00056). The same analyses conducted on the pupal fat bodies also uncovers a similar association between PC1 and nesting locality (Figure S9; *F_1_*_,17_=27.47, p=6.636e-05, R^2^=0.5952).

We then performed a differential expression analysis between these stage-matched pupae from solitary-biased versus social-biased sites (Table S5; Figure S13). We identified 1121 genes differentially expressed in pupal brains (Figure 3d). Genes upregulated in solitary-biased pupal brains are enriched for synaptic transmission and olfactory learning while genes upregulated in social-biased brains are enriched for cell division and protein folding, among others (Figure S13). Of these 1121 genes, 275 are also differentially expressed between adult reproductives and workers, and this overlap is more than expected by chance (hypergeometric test, p=4.49e-23; odds ratio=2.19).

Differences observed between pupae and adults are strongly concordant across social forms (Figure 4). For example, social-biased pupae are predicted to develop into adult workers, and genes upregulated in these pupae overlap significantly with genes upregulated in adult workers. Similarly, solitary-biased pupae are predicted to develop into reproductive adults, and we also observe a significant overlap between genes upregulated in this group and genes upregulated in queens and solitary reproductives. Moreover, the effect size estimates (measured as log2 fold changes) for adult reproductives vs. workers and solitary-biased vs. social-biased pupae are strongly correlated (Figure 4a; linear regression, R^2^=0.41, F=190.24, p=3.38e-33), suggesting that both the magnitude and the directionality of these expression changes are similar in pupal and adult brains. These genes show substantial GO enrichment for chemical synaptic transmission, GABAergic signaling, neurotransmitter receptor metabolism, and synaptic processes, among others (Figure S15). Finally, a random forest classifier demonstrates that pupal brain gene expression is highly predictive of adult behavioral states (Figure S16).

**Figure 4.**
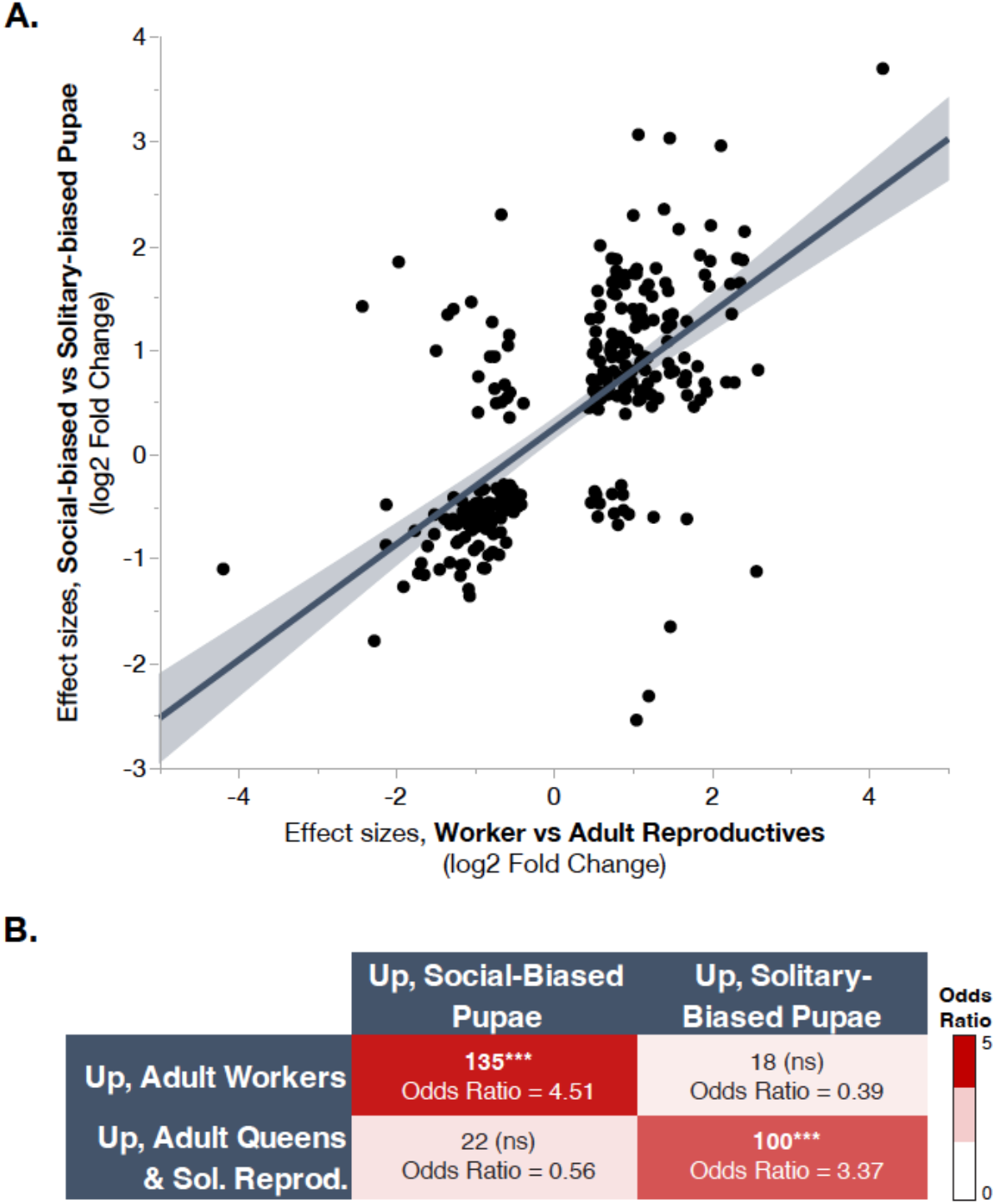
Overlapping DEGs for adult and pupal brain transcriptomes are correlated in direction and magnitude. (A) The effect size estimates (shown here as the log_2_-fold change; DESeq2, FDR<0.001) for overlapping brain DEGs between adult reproductives vs. workers and solitary-biased vs. social-biased pupae are strongly correlated (linear regression, R^2^=0.41, F=190.24, p=3.38e-33). Solid line represents the best fit line for the linear regression, and the shading indicates the 95% confidence intervals around this fit. (B) **DEGs in adults and pupae are highly concordant in expression patterns.** Genes upregulated in social-biased pupal brains are strongly enriched for genes that are also upregulated in adult workers (hypergeometric test, p=8.05e-37, odds ratio=4.51). Similarly, genes more highly expressed in solitary-biased pupae are more likely to also be upregulated in adult queens and solitary reproductives (hypergeometric test, p=1.58e-20, odds ratio=3.37). There is no significant enrichment for discordant patterns of gene expression (e.g., genes upregulated in workers do not also show an enrichment for genes upregulated in solitary-biased pupae, etc.). Asterisks denote a significant enrichment with p<0.0001.

In the fat bodies, we identified 2,028 DEGs between solitary-biased versus social-biased pupae. GO enrichment analyses reveal that genes upregulated in solitary-biased pupae include Toll signaling, cell division, and circadian rhythm, while genes upregulated in social-biased pupae are again enriched for processes linked to cell adhesion and lipid transport, among others (Figure S17). Similar to brains, we found 321 of these DEGs are also differentially expressed in the fat bodies of adult reproductives and workers; this is significantly more than expected by chance (hypergeometric test, p=8.18e-10, odds ratio=1.55). Again, we observe similar concordance across pupal and adult DEGs (Figure S18). As in the brain, we see a significant correlation between effect size estimates (e.g., log2 fold change) across both adult and pupal comparisons (Figure S18a; linear regression, R^2^=0.225, F=92.36, p=2.27e-19).

## Discussion

Here, we explore some of the environmental and molecular factors associated with the variable social strategies employed by *Lasioglossum baleicum.* In this socially flexible species, females can produce either social or solitary nests, and this variation is tightly linked to nesting locality, as supported by previous studies (Cronin and Hirata, 2003; Hirata & Higashi, 2008). Nests constructed in shady patches of the aggregation are largely solitary, while those constructed in sunnier patches are eusocial and contain a reproductive queen and 2-3 nonreproductive workers. This variation is most likely due to behavioral plasticity – there appears to be no genetic divergence between social and solitary nests in this population (Hirata & Higashi, 2008), though more detailed genetic data are needed.

### Spring soil temperature is a strong predictor of social behavior

We found that soil temperatures in early spring, but not in mid-summer, are a strong predictor of the social structure of *L. baleicum* nests. Sites with cooler spring soil temperatures are highly biased towards solitary nests, while sites with warmer spring temperatures are highly biased towards social nests. Summer soil temperatures were not associated with this variation, suggesting that females adjust their nesting strategy depending on early-spring soil conditions at their chosen nest site.

Our results are highly consistent with natural variation in social behavior that has been documented in many social systems (Cronin, 2001; Cronin & Schwarz, 1999; Eickwort et al., 1996; Field et al., 2010; Kocher et al., 2014; Lawson et al., 2018; Plateaux-Quénu, 2008; Purcell, 2011; Rubenstein & Lovette, 2007; Sakagami & Munakata, 1972). Within the social insects, social variation typically occurs across elevational (Sakagami & Munakata, 1972; Soucy & Danforth, 2002) or latitudinal (Cronin & Schwarz, 1999; Davison & Field, 2016; Field et al., 2010) gradients, and solitary nesting strategies are most prevalent when season lengths for foraging and reproduction are shorter.

These geographical patterns in social behavior can be explained, at least in part, by combined constraints in season length as well as the length of the colony reproductive cycle (Kocher et al., 2014). While solitary nests can immediately produce the next generation of reproductives, eusocial nests typically raise non-reproductive workers prior to rearing the next reproductive generation (Michener, 1974; Quiñones & Pen, 2017). In environments with shorter seasons, eusocial nests may simply not have sufficient time to complete reproduction, and season length may therefore act as a strong selective force that shapes variation in social behavior (Kocher et al., 2014; Purcell, 2011). Our findings demonstrate that early spring temperatures can also predict social structure within a local thermal gradient, and these temperature differences may also reflect differences in the length of developmental windows between sunny and shady sites.

### Reproductive status is the strongest predictor of adult gene expression

To better understand the molecular underpinnings of this plasticity, we characterized variation in the gene expression profiles of adult females from solitary and social nests. We find that reproductive status is a strong predictor of gene expression – queens, foundresses, and solitary reproductive all exhibited similar patterns of brain gene expression that were distinct from that of workers. Both social queens and solitary reproductives had highly similar gene expression patterns to each other and to spring foundresses.

Workers in social nests represent the most distinct group in terms of gene expression profiles. Workers do not vary in body size compared to reproductive females, but they do vary in ovarian activation. It is possible that some of these differences could be age-associated because workers have not undergone diapause. However, our mid-season sampling of reproductive females was likely to include both overwintered adult females as well as same-season females from the first brood that either inherited nests as replacement queens or established new nests as solitary females. Thus, the gene expression differences we observed are most likely to be driven by differences in reproductive state rather than behavioral types.

Differentially expressed genes between adult queens and workers were largely associated with reproduction. For example, in queen versus worker brains, the most highly upregulated gene between queens and workers is a gene that encodes an uncharacterized protein predicted to be involved in positive regulation of voltage-gated potassium channel activity and regulation of cholinergic synaptic transmission. In adult fat bodies, *juvenile hormone binding protein 14* (*jhbp14*) is the most strongly upregulated gene in queens compared to workers. Juvenile hormone (JH) is predicted to be involved in various multicellular organism reproduction, and *jhbp14* is involved in the transportation of JH from the *corpora allata* (the glands where JH is synthesized) to all other target cells (Kolodziejczyk et al., 2008). In many insects, JH is a known regulator of development, gonadal activities, diapause, caste determination, reproductive behaviors (Brent et al., 2016; Giray et al., 2005; Nijhout, 1994) and a modulator of sexual receptivity (Bilen et al., 2013; Ringo, 1996; Ringo et al., 1991). Selection on the JH signaling pathway has also been associated with the evolution of social behavior in sweat bees (Jones et al., 2023).

### Pupal gene expression varies across shady, solitary-biased and sunny, social-biased sites

We identified gene expression differences in developing pupae that were associated with variation in environmental conditions among sites. Given the strong correlation between spring soil temperatures and nesting behavior, the gene expression differences we observe are likely to be predictive of future reproductive status: pupae from cooler, solitary-biased nests are predicted to develop into future reproductive females while pupae from warmer, social-biased nests are predicted to develop into non-reproductive workers.

The top DEGs in these groups are largely uncharacterized, but also include genes associated with chitin binding activity and *Osiris 8*. Members of the *Osiris* gene family are known to contribute to phenotypic plasticity in a wide range of social insect species (Smith et al., 2018).

### Pupal expression differences are predictive of adult social forms

We found substantial overlap between the pupal and adult DEGs, and this finding was supported by more rigorous testing using a random forest classifier to demonstrate that gene expression profiles of pupae can be used to predict adult behavior. There is a strong enrichment for concordant expression differences between pupal and adult social forms. For example, social-biased pupae are predicted to develop into adult workers, and genes upregulated in these pupae overlap significantly with genes upregulated in adult workers in both brain and fat body. Similarly, solitary-biased pupae are predicted to develop into reproductive adults, and we observe a significant overlap between genes upregulated in the brains of this group and genes upregulated in queens and solitary reproductives. Interestingly, the concordance between solitary-biased pupae and reproductive adults is less pronounced in the fat bodies for these groups. This difference could potentially be associated with the higher fat stores often observed in newly eclosed adults with high reproductive potential; while developing females grow these fat stores, reproductively active queens and solitary females deplete these fat stores throughout their reproductive period. Finally, for both solitary- and social-biased pupal versus adult comparisons, we find a strong correlation between the effect size estimates of the DEGs in both brains and fat bodies. Taken together, these findings suggest that the gene expression differences in pupae are predictive of adult social behaviors.

Several of the DEGs with concordant gene expression patterns have previously been linked to variation in adult social behavior, but their expression patterns in developing pupae have rarely been characterized. For example, we find that *syntaxin-1a* (*Syx1a*) is upregulated in both social-biased pupae and in adult workers. *Syx1a* is a member of the SNAP/SNARE complex and plays a key role in neurotransmission (Schulze et al., 1995). The expression differences we observe in *L. baleicum* mirror changes in gene expression documented between social and solitary populations of another socially-flexible halictine bee, *Lasioglossum albipes* (Kocher et al., 2018). Our results suggest that changes in *syx1a* expression are not only associated with variation in adult social behaviors in multiple sweat bee species, but that these expression differences may also be relevant during earlier developmental stages. In addition to *syx1a*, we also find concordant changes in gene expression between pupae and adults for a number of genes previously linked to variation in adult social behaviors, including *Tyramine receptor* (*Tyr1*), *NMDA receptor* (*Nmdar1*), *Excitatory amino acid transporter 1* (*Eaat1*), and *Fragile X messenger ribonucleoprotein 1* (*Fmr1*). Each of these genes has been linked to variation in adult social behavior in several social insect systems (Arinrad et al., 2021; Mineur et al., 2006; Spencer et al., 2005; Ujita et al., 2018; Wang et al., 2020). More broadly, these concordant DEGs between pupal and adult brains are strongly enriched for synaptic transmission, long term memory, and GABAergic signaling, suggesting that at least some of the neurobiological changes linked to adult social behaviors may be established earlier during development.

### Developmental environments can shape adult social behaviors in *L. baleicum*

In many organisms, the conditions that occur during early life sensitive periods can generate lasting and sometimes irreversible effects on developmental trajectories (English & Barreaux, 2020). For example, in many social insects, nutritional inequities during early larval development differentiate individuals irreversibly into reproductive queens or non-reproductive workers. Although all adult females in *L. baleicum* are capable of switching between social forms, our results suggest there may also be key differences that take place during *L. baleicum* development that set the stage for adult reproductive status and social behaviors.

Differences in soil temperature could potentially alter the developmental trajectories of offspring in the spring brood. For example, changes in developmental rates could alter expression of different transcriptional networks in developing brood, potentially biasing developmental outcomes towards reproductive states. Warmer temperatures are well known to accelerate development times in many insects (Cui et al., 2018; Van Dis et al 2021). These effects have also been demonstrated directly in *L. baleicum*, where incubation of brood at different temperatures shifts the development times of pupae (Hirata & Higashi, 2008). Similarly, studies of the small carpenter bee, *Ceratina calcarata*, have also demonstrated that warmer developmental environments decrease development times but that they also lead to reduced adult body size and lower survivorship for these individuals (DeHaan et al., 2022).

An additional possibility is that females are differentially provisioning offspring to bias the development of their offspring towards reproductive states (Kapheim et al., 2011). Studies in other sweat bee species have shown that increasing the provision sizes to pupae can alter body size and reproductive status (Lawson et al., 2017; Richards & Packer, 1996). Moreover, in *C. calcarata*, maternal provisions of less nutritious pollen and smaller provision sizes can lead to the production of smaller daughters with less reproductive potential (Lawson et al., 2016). These “dwarf eldest daughters” are more likely to remain in the nest as guards, increasing the fitness benefits to the mother. Whether or not differences in maternal provisions influence caste development in *L. baleicum* remains to be discovered.

Future studies that incorporate finer-scale developmental sampling with environmental manipulations of temperature or diet in sweat bees could provide novel insights into how distinct transcriptional patterns set the stage for distinct behavioral forms in adults.

## Conclusions

Our study provides insights into how environmental factors can shape behavioral variation in a socially flexible sweat bee. Our results reveal that spring soil temperatures are highly predictive of nest social structure and are consistent with theory predicting a trade-off between the reproductive benefits of social nesting versus adequate season lengths required for social reproduction. We find that the primary molecular correlates of adult social behaviors are associated with reproductive status; there are no major differences in the brain gene expression profiles of social versus solitary reproductive females, but workers have a distinct transcriptomic profile. Surprisingly, our results also suggest that the behavioral and reproductive plasticity in this system may not be quite as flexible as previously assumed. We find that developing pupae from social- and solitary-biased nesting sites have distinct transcriptomic profiles, suggesting that the developmental environment may set the stage for adult behaviors even in socially flexible, totipotent species.

## Data Accessibility

The *L. baleicum* genome assembly is available on NCBI, accession GCA_022376115.1. Raw sequence data for all samples have been deposited in NCBI, SRA submission, SUB12095212.

## Author Contributions

KSO, SDK and ALC designed and conceptualized the study. Samples were collected by KSO, SDK, TTHN, ALC and SVG. KSO generated genomic data, and data curation was handled by KSO and AEW. KSO, IMT, and SDK analyzed genomic data. KSO and SDK, generated figures and wrote the initial draft of the manuscript. SDK supervised the work. All authors reviewed and edited the final manuscript.

## Acknowledgements

We thank the Green Management section of the Sapporo City Construction Bureau Green Promotion Department as well as the management of Nishioka Park, Hokkaido, Japan for granting permission to conduct our research, and Scott V. C. Groom for discussion and feedback on field collections. We thank Beryl Jones, Nikki Van Dorp and Weijie Liu for discussion and guidance on molecular protocols, and members of the Kocher Lab for feedback on earlier versions of this manuscript. We also thank the Princeton University Genomic Core staff for sequencing support. This work was supported by grants from the Society for Study of Evolution (SSE) and a Petrie Fellowship to KSO, the Princeton Ogasawara Fund, The Packard Foundation, a Pew Biomedical Scholars Program, NSF DEB 1754476, and NIH 1DP2GM137424-01, the Tokyo Human Resources Fund for City Diplomacy and the Japan Society for the Promotion of Science, Kakenhi 21K06292.

## References

Alexa, A., & Rahnenführer, J. (2007). Gene set enrichment analysis with topGO. Retrieved from http://www.mpi-sb.mpg.de/∼alexa

Arinrad, S., Wilke, J. B. H., Seelbach, A., Doeren, J., Hindermann, M., Butt, U. J., … Ehrenreich, H. (2021). NMDAR1 autoantibodies amplify behavioral phenotypes of genetic white matter inflammation: a mild encephalitis model with neuropsychiatric relevance. Molecular Psychiatry 2021 27:12, 27(12), 4974–4983. https://doi.org/10.1038/s41380-021-01392-8

Benjamini, Y., & Hochberg, Y. (1995). Controlling the False Discovery Rate: A Practical and Powerful Approach to Multiple Testing. Journal of the Royal Statistical Society: Series B (Methodological), 57(1), 289–300. https://doi.org/10.1111/J.2517-6161.1995.TB02031.X

Bilen, J., Atallah, J., Azanchi, R., Levine, J. D., & Riddiford, L. M. (2013). Regulation of onset of female mating and sex pheromone production by juvenile hormone in *Drosophila melanogaster*. Proceedings of the National Academy of Sciences of the United States of America, 110(45), 18321–18326. https://doi.org/10.1073/PNAS.1318119110/-/DCSUPPLEMENTAL

Brand, N., & Chapuisat, M. (2012). Born to be bee, fed to be worker? The caste system of a primitively eusocial insect. Frontiers in Zoology, 9, 35. https://doi.org/10.1186/1742-9994-9-35

Breed, M. D., Silverman, J. M., & Bell, W. J. (1978). Agonistic behavior, social interactions, and behavioral specialization in a primitively eusocial bee. In Insectes Sociaux (Vol. 25).

Brent, C. S., Miyasaki, K., Vuong, C., Miranda, B., Steele, B., Brent, K. G., & Nath, R. (2016). Regulatory roles of biogenic amines and juvenile hormone in the reproductive behavior of the western tarnished plant bug (*Lygus hesperus*). Journal of Comparative Physiology B: Biochemical, Systemic, and Environmental Physiology, 186(2), 169–179. https://doi.org/10.1007/s00360-015-0953-1

Brothers, D. J., & Michener, C. D. (1974). Interactions in colonies of primitively social bees - III. Ethometry of division of labor in *Lasioglossum zephyrum* (Hymenoptera: Halictidae). Journal of Comparative Physiology, 90(2), 129–168. https://doi.org/10.1007/BF00694482/METRICS

Brůna, T., Hoff, K. J., Lomsadze, A., Stanke, M., & Borodovsky, M. (2021). BRAKER2: automatic eukaryotic genome annotation with GeneMark-EP+ and AUGUSTUS supported by a protein database. NAR Genomics and Bioinformatics, 3(1), 1–11. https://doi.org/10.1093/NARGAB/LQAA108

Bryant, D. M., Johnson, K., DiTommaso, T., Tickle, T., Couger, M. B., Payzin-Dogru, D., … Whited, J. L. (2017). A Tissue-Mapped axolotl *de novo* transcriptome enables identification of limb regeneration factors. Cell Reports, 18(3), 762–776. https://doi.org/10.1016/j.celrep.2016.12.063/attachment/b24f06b8-7ac7-45af-b605-0da3b55ae55e/mmc11.zip

Cantarel, B. L., Korf, I., Robb, S. M. C., Parra, G., Ross, E., Moore, B., … Yandell, M. (2008). MAKER: An easy-to-use annotation pipeline designed for emerging model organism genomes. Genome Research, 18(1), 188–196. https://doi.org/10.1101/GR.6743907

Chen, S., Zhou, Y., Chen, Y., & Gu, J. (2018). fastp: an ultra-fast all-in-one FASTQ preprocessor. Bioinformatics, 34(17), i884–i890. https://doi.org/10.1093/BIOINFORMATICS/BTY560

Conway, J. R., Lex, A., & Gehlenborg, N. (2017). UpSetR: an R package for the visualization of intersecting sets and their properties. Bioinformatics, 33(18), 2938–2940. https://doi.org/10.1093/bioinformatics/btx364

Cronin, A.L., & Hirata, and M. (2003). Social polymorphism in the sweat bee *Lasioglossum* (*Evylaeus*) *baleicum* (Cockerell) (Hymenoptera, Halictidae) in Hokkaido, northern Japan. Insectes Sociaux, 379–386. https://doi.org/10.1007/s00040-003-0693-1

Cronin, Adam L. (2001). Social flexibility in a primitively social allodapine bee (Hymenoptera: Apidae): Results of a translocation experiment. Oikos, 94(2), 337–343. https://doi.org/10.1034/j.1600-0706.2001.940214.x

Cronin, Adam L., & Schwarz, M. P. (1999a). Latitudinal variation in the life cycle of allodapine bees (Hymenoptera; Apidae). Canadian Journal of Zoology, 77(6), 857–864. https://doi.org/10.1139/CJZ-77-6-857

Cronin, Adam L., & Schwarz, M. P. (1999b). Life cycle and social behavior in a heathland population of *Exoneura robusta* (Hymenoptera: Apidae): habitat influences opportunities for sib rearing in a primitively social bee. Annals of the Entomological Society of America, 92(5), 707–716. https://doi.org/10.1093/AESA/92.5.707

Cui, J., Zhu, S. Y., Bi, R., Xu, W., Gao, Y., & Shi, S. Sen. (2018). Effect of temperature on the development, survival, and fecundity of *Heliothis viriplaca* (Lepidoptera: Noctuidae). Journal of Economic Entomology, 111(4), 1940–1946. https://doi.org/10.1093/JEE/TOY151

Davison, P. J., & Field, J. (2016). Social polymorphism in the sweat bee *Lasioglossum* (*Evylaeus*) *calceatum*. Insectes Sociaux, 63(2), 327–338. https://doi.org/10.1007/s00040-016-0473-3

Davison, P. J., & Field, J. (2018). Limited social plasticity in the socially polymorphic sweat bee *Lasioglossum calceatum*. Behavioral Ecology and Sociobiology, 72(3). https://doi.org/10.1007/s00265-018-2475-9

Dehaan, J. L., Maretzki, J., Skandalis, A., Tattersall, G. J., & Richards, M. H. (2022). Costs and benefits of maternal nest choice: tradeoffs between brood survival and thermal stress for small carpenter bees. https://doi.org/10.1101/2022.11.30.518597

Dobin, A., Davis, C. A., Schlesinger, F., Drenkow, J., Zaleski, C., Jha, S., … Gingeras, T. R. (2013). STAR: Ultrafast universal RNA-seq aligner. Bioinformatics, 29(1), 15–21. https://doi.org/10.1093/bioinformatics/bts635

Eickwort, G. C., Eickwort, J. M., Gordon, J., Eickwort, M. A., & Weislo, W. T. (1996). Solitary behavior in a high-altitude population of the social sweat bee *Halictus rubicundus* (Hymenoptera: Halictidae). Behavioral Ecology and Sociobiology, 38(4), 227–233. https://doi.org/10.1007/s002650050236

English, S., & Barreaux, A. M. (2020). The evolution of sensitive periods in development: insights from insects. Current Opinion in Behavioral Sciences, 36, 71–78. https://doi.org/10.1016/J.COBEHA.2020.07.009

Field, J., Paxton, R. J., Soro, A., & Bridge, C. (2010). Cryptic plasticity underlies a major evolutionary transition. Current Biology, 20(22), 2028–2031. https://doi.org/10.1016/j.cub.2010.10.020

Field, J., Paxton, R., Soro, A., Craze, P., & Bridge, C. (2012). Body size, demography and foraging in a socially plastic sweat bee: A common garden experiment. Behavioral Ecology and Sociobiology. https://doi.org/10.1007/s00265-012-1322-7

Gabriel, L., Hoff, K. J., Brůna, T., Borodovsky, M., & Stanke, M. (2021). TSEBRA: transcript selector for BRAKER. BMC Bioinformatics, 22(1), 1–12. https://doi.org/10.1186/S12859-021-04482-0/FIGURES/3

Giray, T., Giovanetti, M., & West-Eberhard, M. J. (2005). Juvenile hormone, reproduction, and worker behavior in the neotropical social wasp *Polistes canadensis*. Proceedings of the National Academy of Sciences of the United States of America, 102(9), 3330–3335. https://doi.org/10.1073/pnas.0409560102

Grabherr, M. G., Haas, B. J., Yassour, M., Levin, J. Z., Thompson, D. A., Amit, I., … Regev, A. (2011). Full-length transcriptome assembly from RNA-Seq data without a reference genome. Nature Biotechnology 2011 29:7, 29(7), 644–652. https://doi.org/10.1038/nbt.1883

Haas, B. J., Delcher, A. L., Mount, S. M., Wortman, J. R., Smith, R. K., Hannick, L. I., … White, O. (2003). Improving the *Arabidopsis* genome annotation using maximal transcript alignment assemblies. Nucleic Acids Research, 31(19), 5654–5666. https://doi.org/10.1093/NAR/GKG770

Hirata, M., & Higashi, S. (2008). Degree-day accumulation controlling allopatric and sympatric variations in the sociality of sweat bees, *Lasioglossum* (*Evylaeus*) *baleicum* (Hymenoptera: Halictidae). Behavioral Ecology and Sociobiology, 62(8), 1239–1247. https://doi.org/10.1007/s00265-008-0552-1

Huang, N., & Li, H. (2023). miniBUSCO: a faster and more accurate reimplementation of BUSCO. BioRxiv, 2023.06.03.543588. https://doi.org/10.1101/2023.06.03.543588

Jones, B. M., Rubin, B. E. R., Dudchenko, O., Kingwell, C. J., Traniello, I. M., Wang, Z. Y., … Kocher, S. D. (2023). Convergent and complementary selection shaped gains and losses of eusociality in sweat bees. Nature Ecology and Evolution. https://doi.org/10.1038/s41559-023-02001-3

Jones, P., Binns, D., Chang, H. Y., Fraser, M., Li, W., McAnulla, C., … Hunter, S. (2014). InterProScan 5: genome-scale protein function classification. Bioinformatics, 30(9), 1236–1240. https://doi.org/10.1093/BIOINFORMATICS/BTU031

Kapheim, K. M., Bernal, S. P., Smith, A. R., Nonacs, P., & Wcislo, W. T. (2011). Support for maternal manipulation of developmental nutrition in a facultatively eusocial bee, *Megalopta genalis* (Halictidae). Behavioral Ecology and Sociobiology, 65(6), 1179–1190. https://doi.org/10.1007/S00265-010-1131-9/TABLES/3

Kapheim, K. M., Smith, A. R., Ihle, K. E., Amdam, G. V., Nonacs, P., & Wcislo, W. T. (2012). Physiological variation as a mechanism for developmental caste-biasing in a facultatively eusocial sweat bee. Proceedings of the Royal Society B: Biological Sciences, 279(1732), 1437–1446. https://doi.org/10.1098/RSPB.2011.1652

Kevin blighe. (2022). EnhancedVolcano: Publication-ready volcano plots with enhanced colouring and labeling. Retrieved October 1, 2022, from https://github.com/kevinblighe/EnhancedVolcano

Kocher, S. D., Mallarino, R., Rubin, B. E. R., Yu, D. W., Hoekstra, H. E., & Pierce, N. E. (2018). The genetic basis of a social polymorphism in halictid bees. Nature Communications, 9(1). https://doi.org/10.1038/s41467-018-06824-8

Kocher, S. D., Pellissier, L., Veller, C., Purcell, J., Nowak, M. A., Chapuisat, M., & Pierce, N. E. (2014). Transitions in social complexity along elevational gradients reveal a combined impact of season length and development time on social evolution. Proceedings of the Royal Society B: Biological Sciences, 281(1787). https://doi.org/10.1098/rspb.2014.0627

Kolodziejczyk, R., Bujacz, G., Jakób, M., Ozyhar, A., Jaskolski, M., & Kochman, M. (2008). Insect juvenile hormone binding protein shows ancestral fold present in human lipid-binding proteins. Journal of Molecular Biology, 377(3), 870–881. https://doi.org/10.1016/j.jmb.2008.01.026

Kriventseva, E. V., Kuznetsov, D., Tegenfeldt, F., Manni, M., Dias, R., Simão, F. A., & Zdobnov, E. M. (2019). OrthoDB v10: sampling the diversity of animal, plant, fungal, protist, bacterial and viral genomes for evolutionary and functional annotations of orthologs. Nucleic Acids Research, 47(D1), D807–D811. https://doi.org/10.1093/NAR/GKY1053

Lawson, S P, Shell, W. A., Lombard, · S S, & Rehan, · S M. (2018). Climatic variation across a latitudinal gradient affect phenology and group size, but not social complexity in small carpenter bees. Insectes Sociaux, 65, 483–492. https://doi.org/10.1007/s00040-018-0635-6

Lawson, Sarah P., Helmreich, S. L., & Rehan, S. M. (2017). Effects of nutritional deprivation on development and behavior in the subsocial bee *Ceratina calcarata* (Hymenoptera: Xylocopinae). Journal of Experimental Biology, 220(23), 4456–4462. https://doi.org/10.1242/jeb.160531

Lawson, Sarah P, Ciaccio, K. N., & Rehan, S. M. (2016). Maternal manipulation of pollen provisions affects worker production in a small carpenter bee. Behavioral Ecology and Sociobiology. https://doi.org/10.1007/s00265-016-2194-z

Leek, J. T., & Storey, J. D. (2007). Capturing heterogeneity in gene expression studies by surrogate variable analysis. PLOS Genetics, 3(9), e161. https://doi.org/10.1371/JOURNAL.PGEN.0030161

Liao, Y., Smyth, G. K., & Shi, W. (2014). featureCounts: an efficient general purpose program for assigning sequence reads to genomic features. Bioinformatics, 30(7), 923–930. https://doi.org/10.1093/BIOINFORMATICS/BTT656

Liaw, A., & Wiener, M. (2002). Classification and Regression by randomForest. 2(3). Retrieved from http://www.stat.berkeley.edu/

Love, M. I., Huber, W., & Anders, S. (2014). Moderated estimation of fold change and dispersion for RNA-seq data with DESeq2. Genome Biology, 15(12), 550. https://doi.org/10.1186/s13059-014-0550-8

Michener, C. D. (1974). The Social Behavior of the Bees. Harvard University Press, Cambridge, MA.

Mineur, Y. S., Huynh, L. X., & Crusio, W. E. (2006). Social behavior deficits in the Fmr1 mutant mouse. Behavioural Brain Research, 168(1), 172–175. https://doi.org/10.1016/J.BBR.2005.11.004

Nijhout, H. F. (1994). Insect hormones. Princeton University Press, Princeton, NJ.

Perry, M. (2022). heatmaps: Flexible Heatmaps for Functional Genomics and Sequence Features. R package version 1.20.0. Retrieved October 15, 2022, from https://bioconductor.org/packages/release/bioc/html/heatmaps.html

Plateaux-Quénu, C. (2008). Subsociality in halictine bees. Insectes Sociaux, 55(4), 335–346. https://doi.org/10.1007/S00040-008-1028-Z/METRICS

Plateaux-Quénu, C., Plateaux, L., & Packer, L. (2000). Population-typical behaviours are retained when eusocial and non-eusocial forms of *Evylaeus albipes* (F.) (Hymenoptera, Halictidae) are reared simultaneously in the laboratory. Insectes Sociaux, 47(3), 263–270. https://doi.org/10.1007/PL00001713

Purcell, J. (2011). Geographic patterns in the distribution of social systems in terrestrial arthropods. Biological Reviews, 86(2), 475–491. https://doi.org/10.1111/J.1469-185X.2010.00156.X

Quiñones, A. E., & Pen, I. (2017). A unified model of Hymenopteran preadaptations that trigger the evolutionary transition to eusociality. Nature Communications 2017 8:1, 8(1), 1–13. https://doi.org/10.1038/ncomms15920

Renaud, G., Stenzel, U., Maricic, T., Wiebe, V., & Kelso, J. (2015). deML: robust demultiplexing of Illumina sequences using a likelihood-based approach. Bioinformatics (Oxford, England), 31(5), 770–772. https://doi.org/10.1093/BIOINFORMATICS/BTU719

Richards, M. H., & Packer, L. (1996). The socioecology of body size variation in the primitively eusocial sweat bee, *Halictus ligatus* (Hymenoptera: Halictidae). Oikos, 77(1), 68. https://doi.org/10.2307/3545586

Ringo, J. (1996). Sexual receptivity in insects. Annual Review of Entomology, 41, 473–494. Retrieved from www.annualreviews.org

Ringo, J., Werczberger, R., Altaratz, M., & Segal, D. (1991). Female sexual receptivity is defective in juvenile hormone-deficient mutants of the *apterous* gene of *Drosophila melanogaster*. Behavior Genetics, 21(5), 453–469. https://doi.org/10.1007/BF01066724

Rubenstein, D. R., & Lovette, I. J. (2007). Temporal environmental variability drives the evolution of cooperative breeding in birds. Current Biology, 17(16), 1414–1419. https://doi.org/10.1016/J.CUB.2007.07.032

Sakagami, S. F., & Munakata, M. (1972). Distribution and bionomics of a transpalaearctic eusocial halictine bee, Lasioglossum (Evylaeus) calceatum, in northern Japan, with reference to its solitary life cycle at high altitude. 北海道大學理學部紀要, 18(3), 411–439.

Saleh, N. W., & Ramírez, S. R. (2019). Sociality emerges from solitary behaviours and reproductive plasticity in the orchid bee *Euglossa dilemma*. Royal Society Publishing. https://doi.org/10.1098/rspb.2019.0588

Schulze, K. L., Broadie, K., Perin, M. S., & Bellen, H. J. (1995). Genetic and electrophysiological studies of *Drosophila* syntaxin-1A demonstrate its role in nonneuronal secretion and neurotransmission. Cell, 80(2), 311–320. https://doi.org/10.1016/0092-8674(95)90414-X

Schwarz, M. P., Richards, M. H., & Danforth, B. N. (2007). Changing paradigms in insect social evolution: Insights from halictine and allodapine Bees. Annual Review of Entomology, 52(1), 127–150. https://doi.org/10.1146/annurev.ento.51.110104.150950

Sheehan, M. J., Botero, C. A., Hendry, T. A., Sedio, B. E., Jandt, J. M., Weiner, S., … Tibbetts, E. A. (2015). Different axes of environmental variation explain the presence vs. extent of cooperative nest founding associations in *Polistes* paper wasps. Ecology Letters, 18(10), 1057–1067. https://doi.org/10.1111/ELE.12488

Simão, F. A., Waterhouse, R. M., Ioannidis, P., Kriventseva, E. V., & Zdobnov, E. M. (2015). BUSCO: assessing genome assembly and annotation completeness with single-copy orthologs. Bioinformatics, 31(19), 3210–3212. https://doi.org/10.1093/BIOINFORMATICS/BTV351

Smith, C. R., Morandin, C., Noureddine, M., & Pant, S. (2018). Conserved roles of *Osiris* genes in insect development, polymorphism and protection. Journal of Evolutionary Biology, 31(4), 516–529. https://doi.org/10.1111/JEB.13238

Soucy, S. L., & Danforth, B. N. (2002). Phylogeography of the socially polymorphic sweat bee *Halictus rubicundus* (hymenoptera: Halictidae). Evolution, 56(2), 330–341. https://doi.org/10.1111/j.0014-3820.2002.tb01343.x

Spencer, C. M., Alekseyenko, O., Serysheva, E., Yuva-Paylor, L. A., & Paylor, R. (2005). Altered anxiety-related and social behaviors in the *Fmr1* knockout mouse model of fragile X syndrome. *Genes*, Brain and Behavior, 4(7), 420–430. https://doi.org/10.1111/J.1601-183X.2005.00123.X

Steitz, I., & Ayasse, M. (2020). Macrocyclic lactones act as a queen pheromone in a primitively eusocial sweat bee. Current Biology. https://doi.org/10.1016/j.cub.2020.01.026

Tian, L., & Hines, H. M. (2018). Morphological characterization and staging of bumble bee pupae. PeerJ, 2018(12). https://doi.org/10.7717/PEERJ.6089/SUPP-6

Tom, T., Tom, T., Brůna, T., Brůna, B., Lomsadze, A., & Borodovsky, M. (2020). GeneMark-EP+: eukaryotic gene prediction with self-training in the space of genes and proteins. NAR Genomics and Bioinformatics, 2(2). https://doi.org/10.1093/NARGAB/LQAA026

Ujita, W., Kohyama-Koganeya, A., Endo, N., Saito, T., & Oyama, H. (2018). Mice lacking a functional NMDA receptor exhibit social subordination in a group-housed environment. The FEBS Journal, 285(1), 188–196. https://doi.org/10.1111/FEBS.14334

Van Dis, N. E., Van Der Zee, M., Hut, R. A., Wertheim, B., & Visser, M. E. (2021). Timing of increased temperature sensitivity coincides with nervous system development in winter moth embryos. The Journal of Experimental Biology, 224(17). https://doi.org/10.1242/JEB.242554

Wang, Y., Amdam, G. V., Daniels, B. C., & Page, R. E. (2020). Tyramine and its receptor TYR1 linked behavior QTL to reproductive physiology in honey bee workers (*Apis mellifera*). Journal of Insect Physiology, 126, 104093. https://doi.org/10.1016/J.JINSPHYS.2020.104093

Weisenfeld, N. I., Kumar, V., Shah, P., Church, D. M., & Jaffe, D. B. (2017). Direct determination of diploid genome sequences. Genome Research, 27(5), 757–767. https://doi.org/10.1101/GR.214874.116

Yagi, N., & Hasegawa, E. (2011). Social-organization shift in the sweat bee, *Lasioglossum baleicum* (Hymenoptera, Halictidae), corresponds to changes in foraging activity of the predatory ant Tetramorium tsushimae (Hymenoptera, Formicidae). Sociobiology, 58(1), 241–250.

Yagi, N., & Hasegawa, E. (2012). A halictid bee with sympatric solitary and eusocial nests offers evidence for Hamilton’s rule. Nature Communications. https://doi.org/10.1038/ncomms1939

