## Supplementary information for "Developmental transcriptomes predict adult social behaviors in the socially flexible sweat bee, *Lasioglossum baleicum*"

### Supplementary Data Files

**Table S1.** Sample information and metadata

**Table S2.** Genome assembly statistics.

**Table S3.** Raw gene counts for each sample

**Figure S4.** Intertegular distance across social forms

**Table S5.** Differentially expressed genes for all contrasts

**Figure S6.** Heatmaps and sample clustering, adults

**Figure S7.** Upset plots, adult brains

**Figure S8.** Upset plots, adult fat bodies

**Figure S9.** Principal component analysis and volcano plots, fat bodies

**Table S10.** Gene Ontology analysis results for all contrasts

**Figure S11.** Significantly enriched GO terms, adult reproductives vs workers, brains

**Figure S12.** Significantly enriched GO terms, adult reproductives vs workers, fat bodies

**Figure S13.** Heatmaps and sample clustering, pupae

**Figure S14.** Significantly enriched GO terms, social-biased vs solitary-biased pupal, brains

**Figure S15.** TopGO concordant DEGs, brains

**Figure S16.** Random forest classification of adult behavioral state using pupal brain differentially expressed genes

**Figure S17.** Significantly enriched GO terms, social-biased vs solitary-biased pupal, fat bodies

**Figure S18.** Overlapping DEGs for adult and pupal fat body transcriptomes

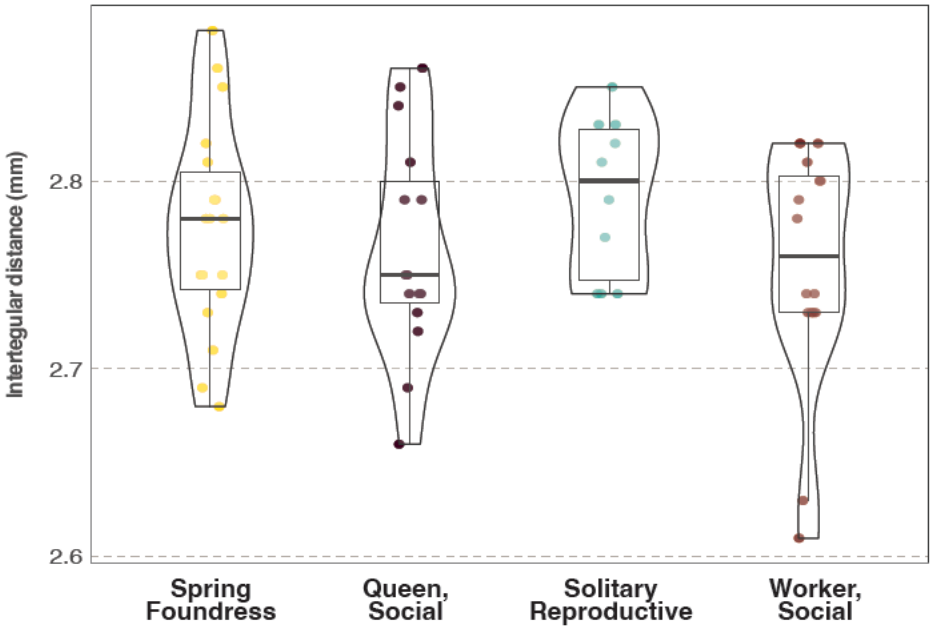


**Figure S4. Intertegular distance does not vary across social forms.** Morphological analysis of all the adults intertegular distance (Kruskal-Wallis: $X_3^2 = 3.47$, $p$ =0.32). N = 62 (Queens = 15, Foundress = 19, solitary reproductives = 11, workers = 17).

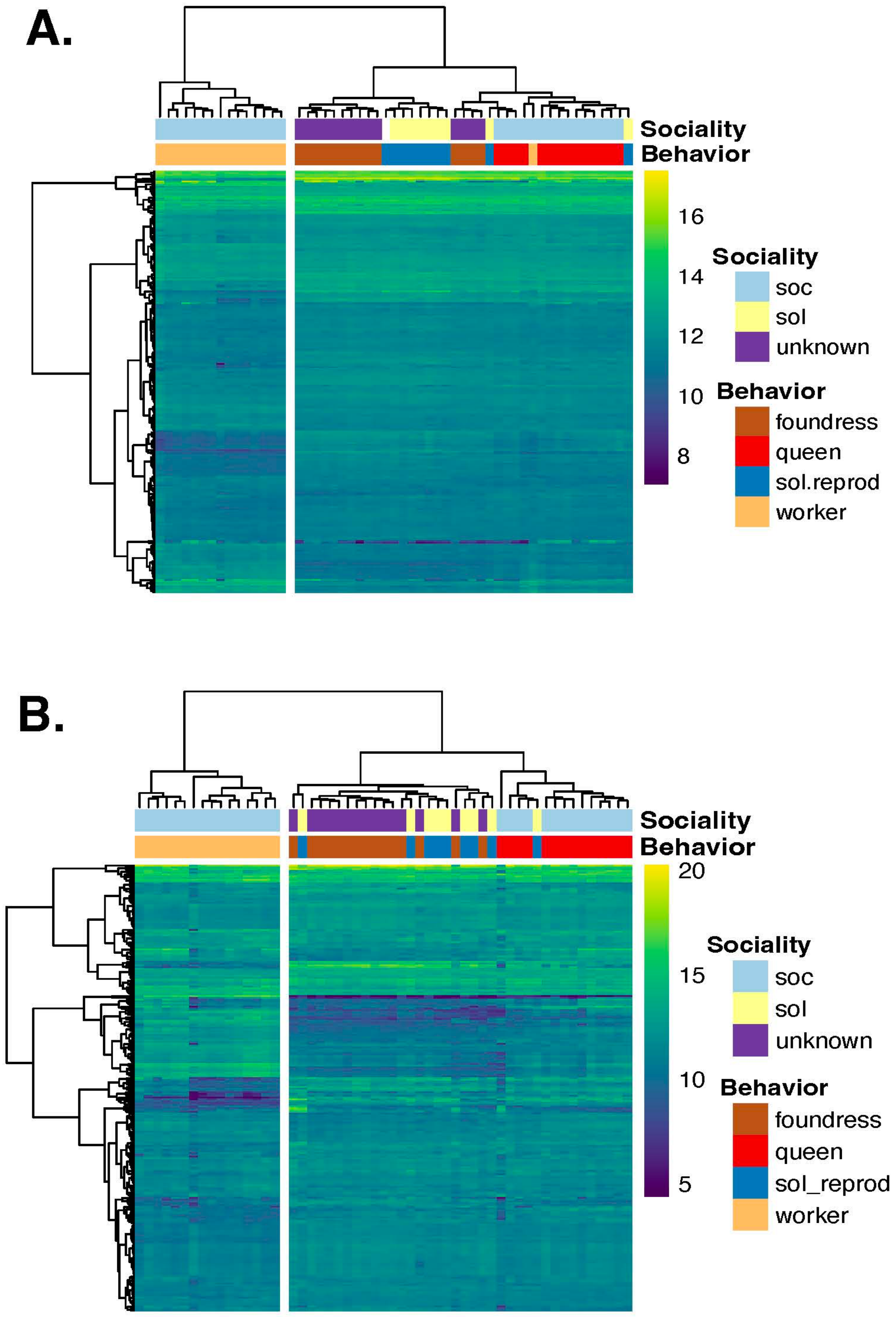


**Figure S6. Hierarchical clustering of adult brain and fat body transcriptomes.** Heatmap of top 250 clustered adult genes in A) brains and B) fat body genes. Scale bar indicates values of variance transformed read counts with *vst()* function in DESeq2 package on R. At the top, the two main branches from the dendrogram trees present grouping by the reproduction role of the individuals. On the left are non-reproductive workers while reproductive queens, solitary reproductives and foundresses form a bigger cluster to the right.

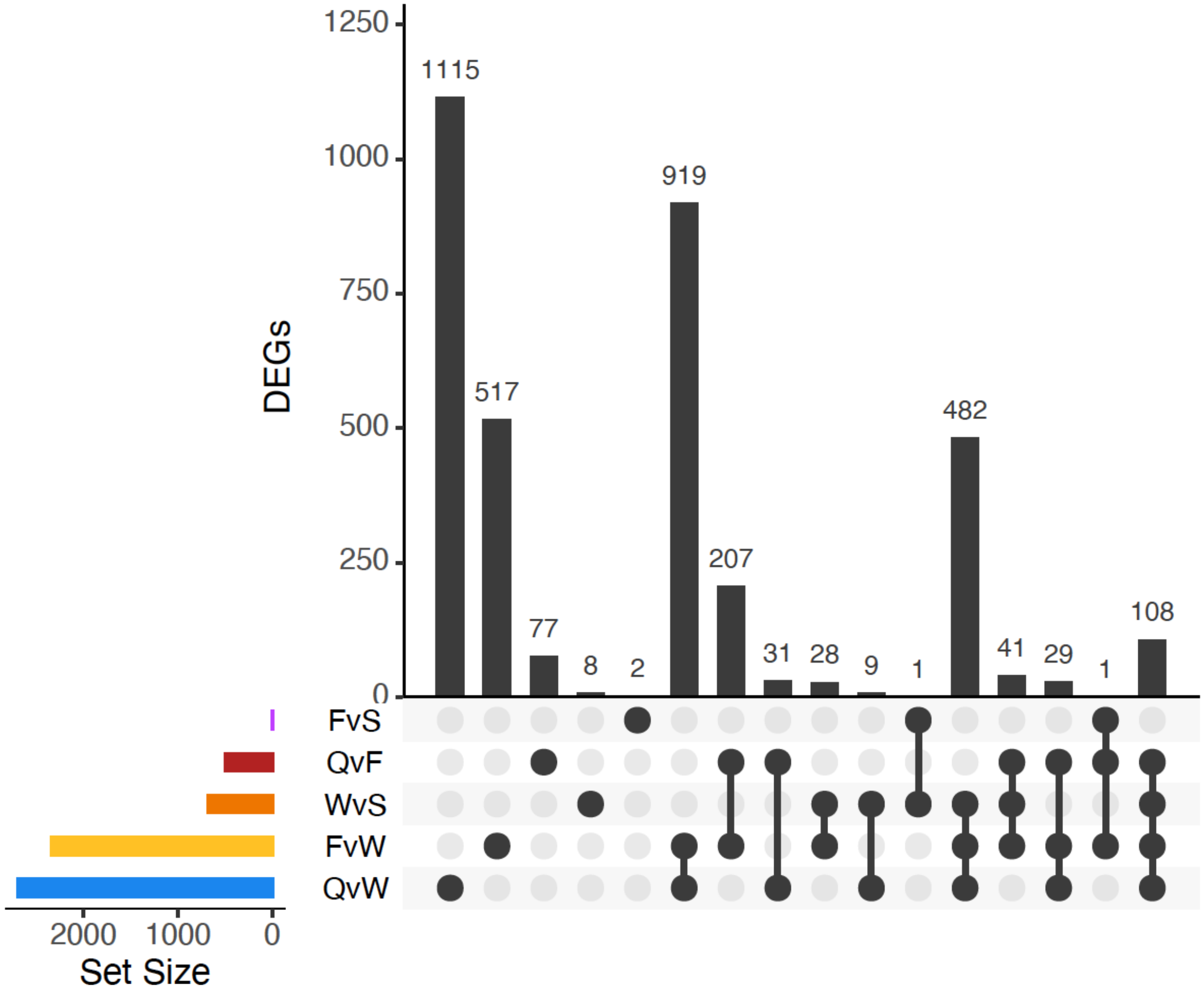


**Figure S7.** Upset plots comparing overlap among DEGs for all pairwise comparisons of adult females, brain. Set size is the number of DEGs identified for each pairwise comparison. F= foundress; S=Solitary Reproductive; W=Workers, Q=Queens.

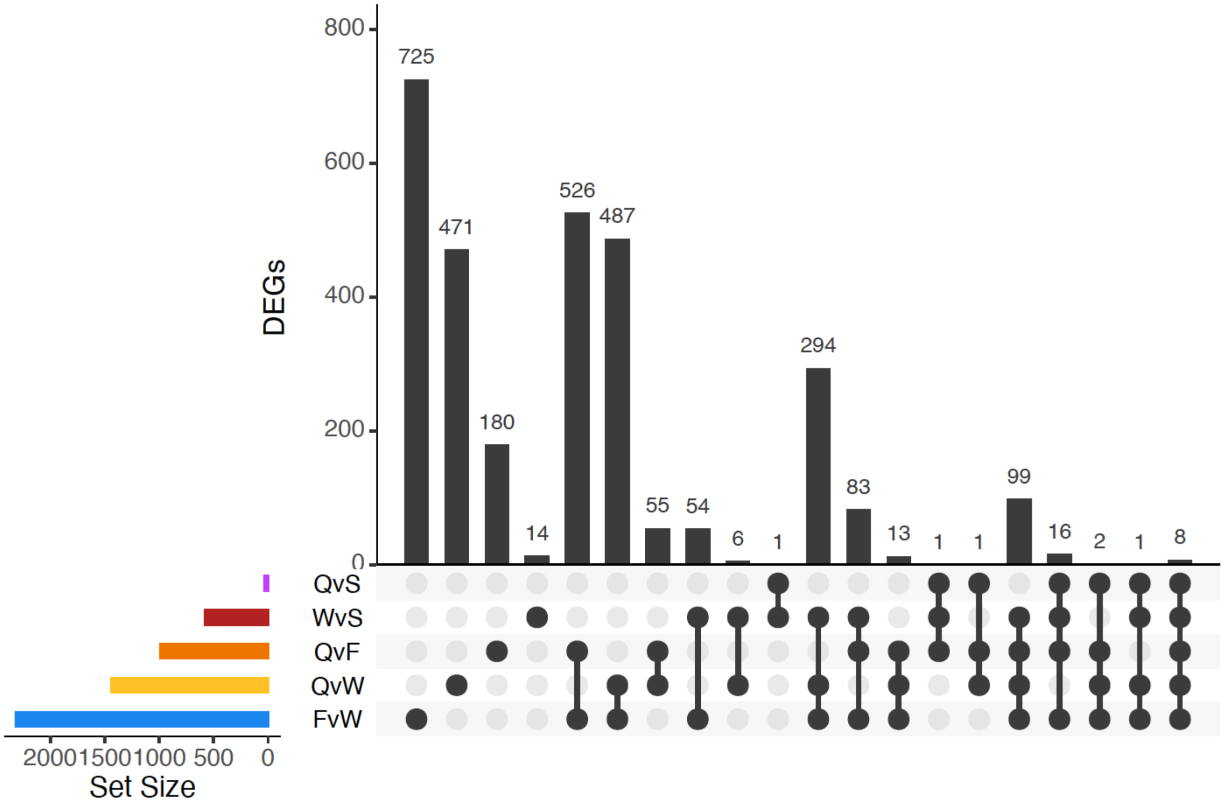


**Figure S8.** Upset plots comparing overlap among DEGs for all pairwise-comparisons of adult females, fat body. Set size is the number of DEGs identified for each pairwise comparison. F= foundress; S=Solitary Reproductive; W=Workers, Q=Queens.

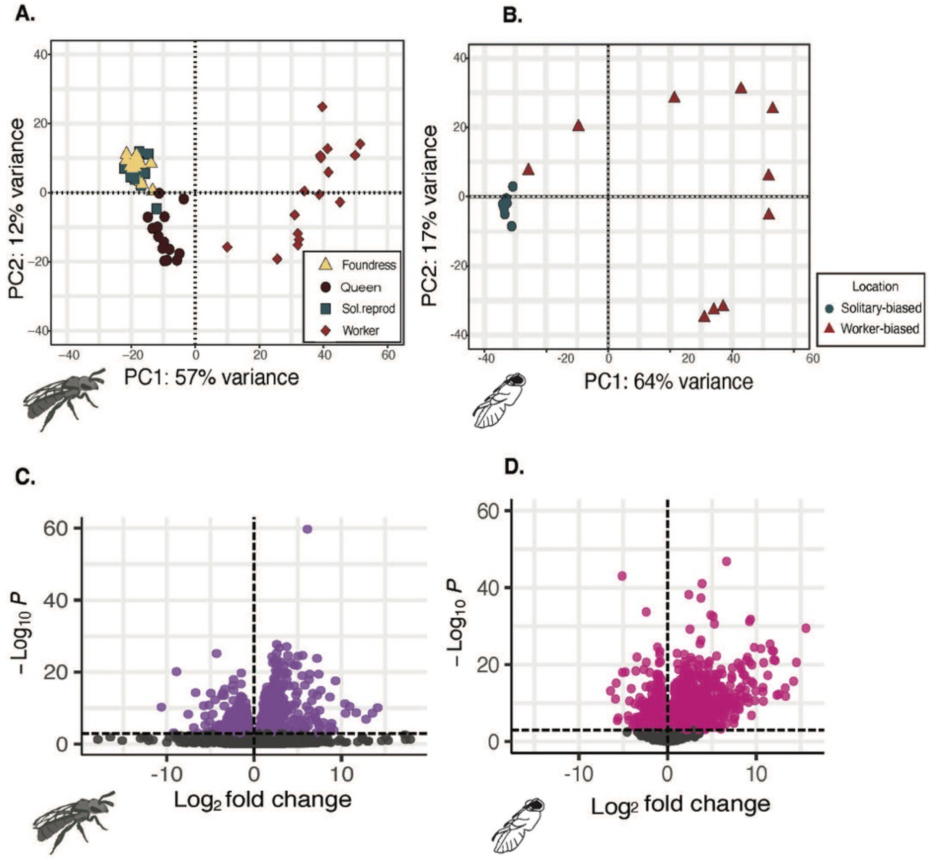


**Figure S9. Adult and pupal fat body gene expression patterns reflect environmental conditions and social behavior.** PCA on all fat body transcripts discriminates between adult behavioral forms (A) as well between late spring pupae excavated from eusocial-biased (sunny) and solitary-biased (shady) sites. Pupae from sunny sites are most likely worker-destined, while pupae excavated from shady sites are predicted to become reproductive females. (C-D). Volcano plots highlighting differentially expressed genes (DEGs). (C) reproductive adults (queens and solitary reproductives) vs workers (n = 1221 DEGs). DEG's were all called using DEseq2 (FDR <0.001). (D) Reproductive-biased pupae from the shaded sites versus worker-biased pupae from sites with full sun (n=2028 DEGs).

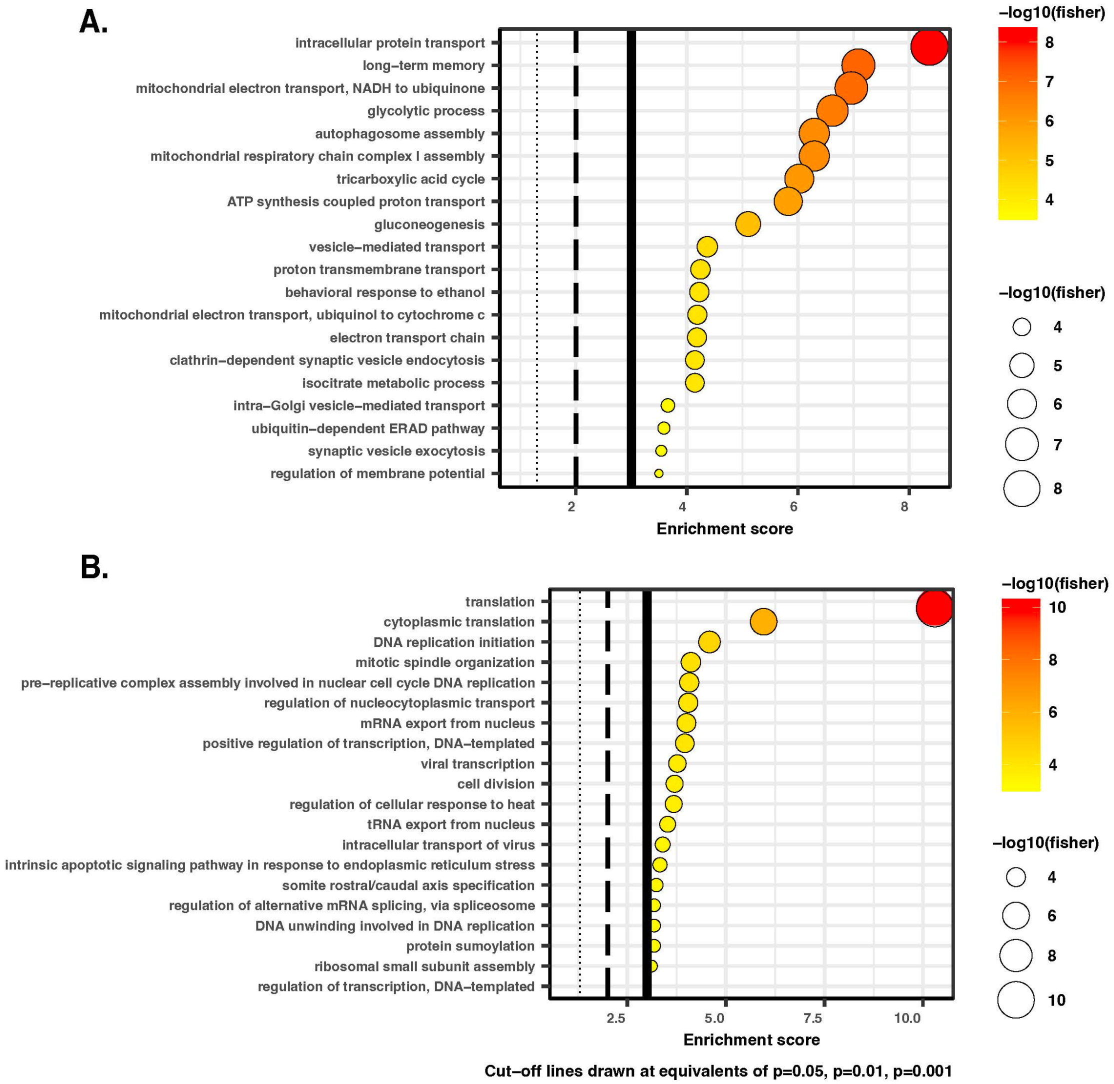


**Figure S11. Significant enrichment for GO Biological Processes, adult brains.** A. GO enrichments for brain DEGs upregulated in workers vs adult queens and solitary reproductives. B. GO enrichments for brain DEGs downregulated in adult queens and solitary reproductives compared to workers. Results are visualized using TopGO. Circle diameter and color is proportional to a -log10 pvalue from a Fisher Exact Test for enrichment of each GO term. Three significance threshold lines are included, for $p<0.05$ (dotted), $p<0.01$ (dashed), and $p<0.001$ (solid).

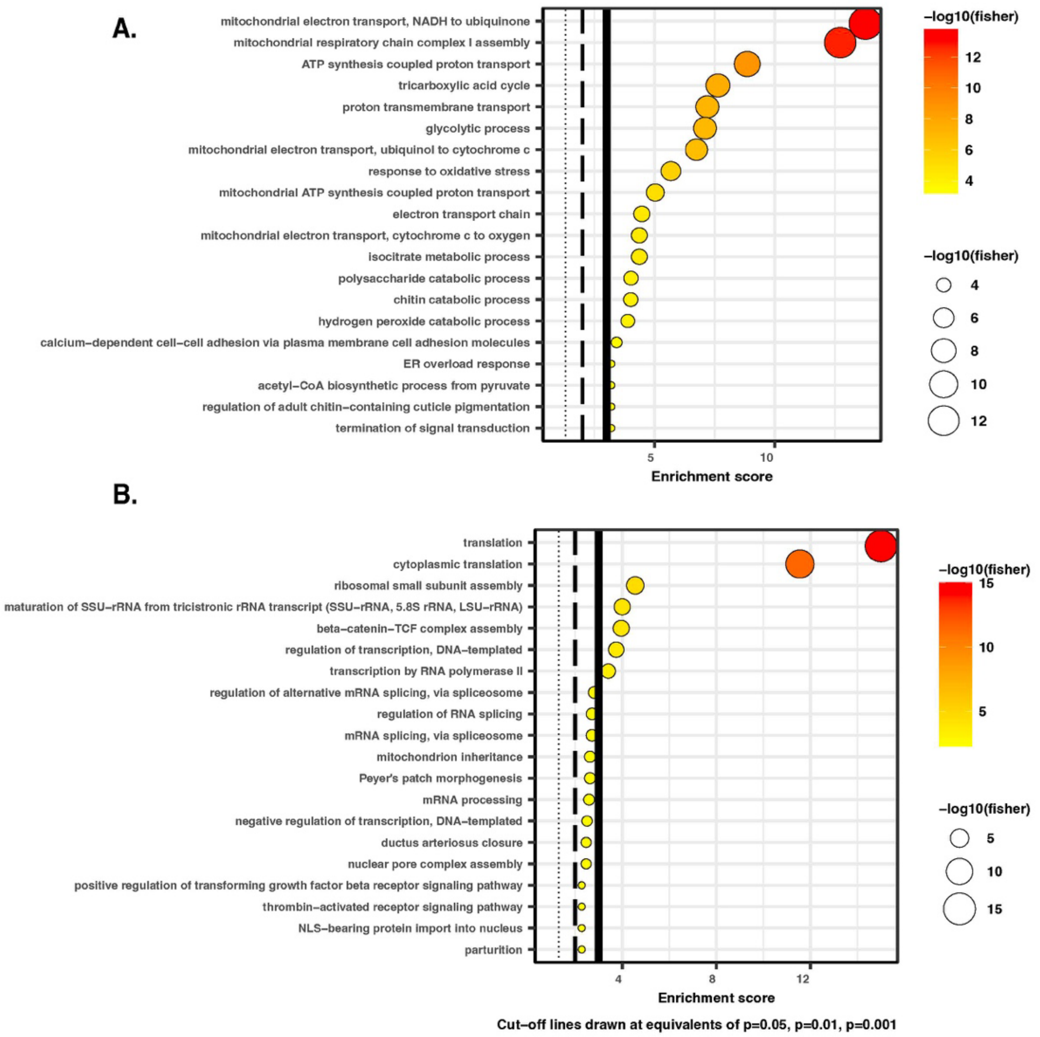


**Figure S12**. **Significant enrichment for GO Biological Processes, adult fatbodies**.. A. GO enrichments for fat body DEGs upregulated in adult queens and solitary reproductives compared to workers. B. GO enrichments for fat body DEGs downregulated in adult queens and solitary reproductives compared to workers. Results are visualized using TopGO. Circle diameter and color is proportional to a -log10 pvalue from a Fisher Exact Test for enrichment of each GO term. Three significance threshold lines are included, for $p<0.05$ (dotted), $p<0.01$ (dashed), and $p<0.001$ (solid).

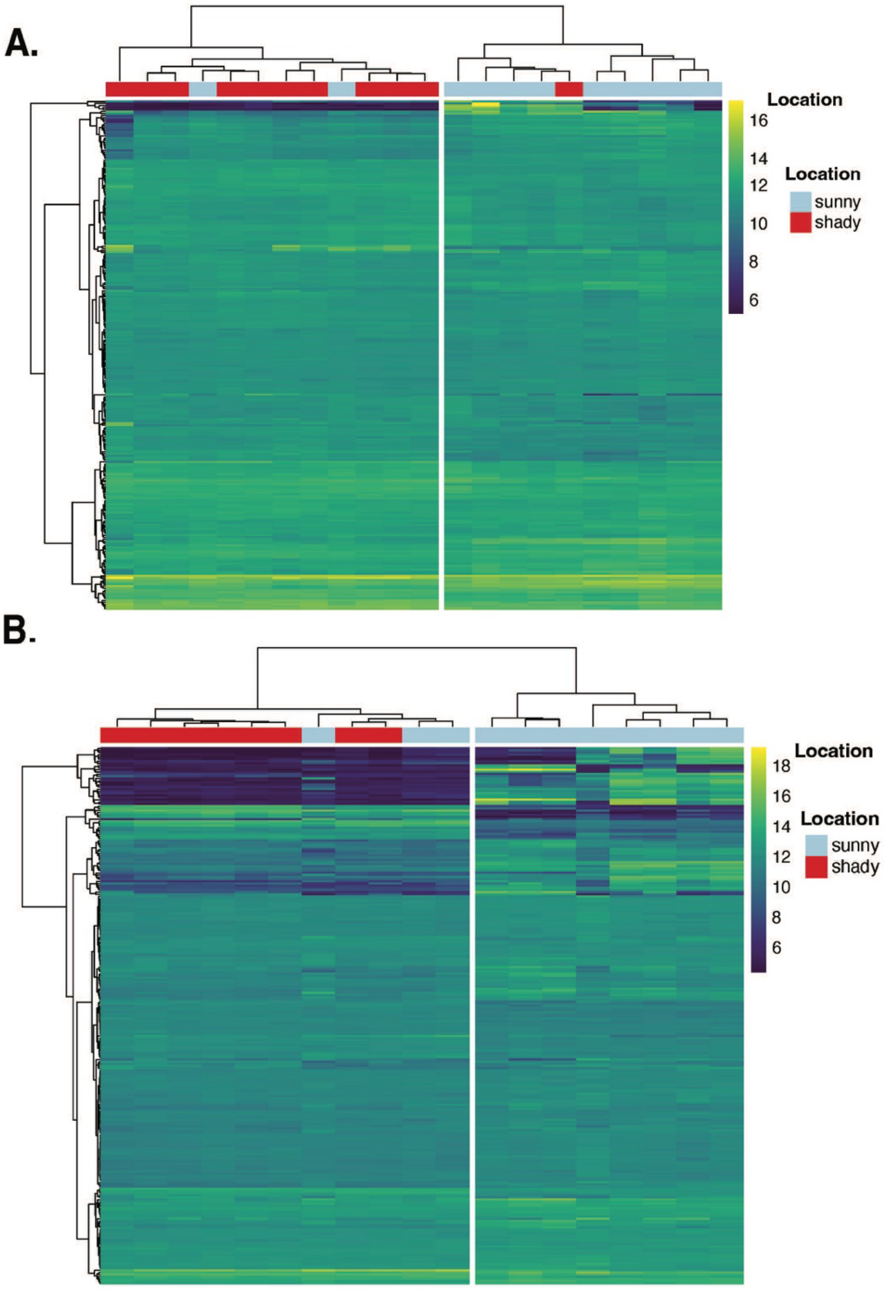


**Figure S13. Hierarchical clustering of pupal brain and fat body transcriptomes.** Heatmap of top 250 clustered pupal genes in A) brains and B) fat body genes. Scale bar indicates values of variance transformed read counts with *vst()* function in DESeq2 package in R. At the top, the two main branches from the dendrogram trees present grouping by nest locations. On the left are social-biased pupa while solitary-biased pupa cluster to the right.

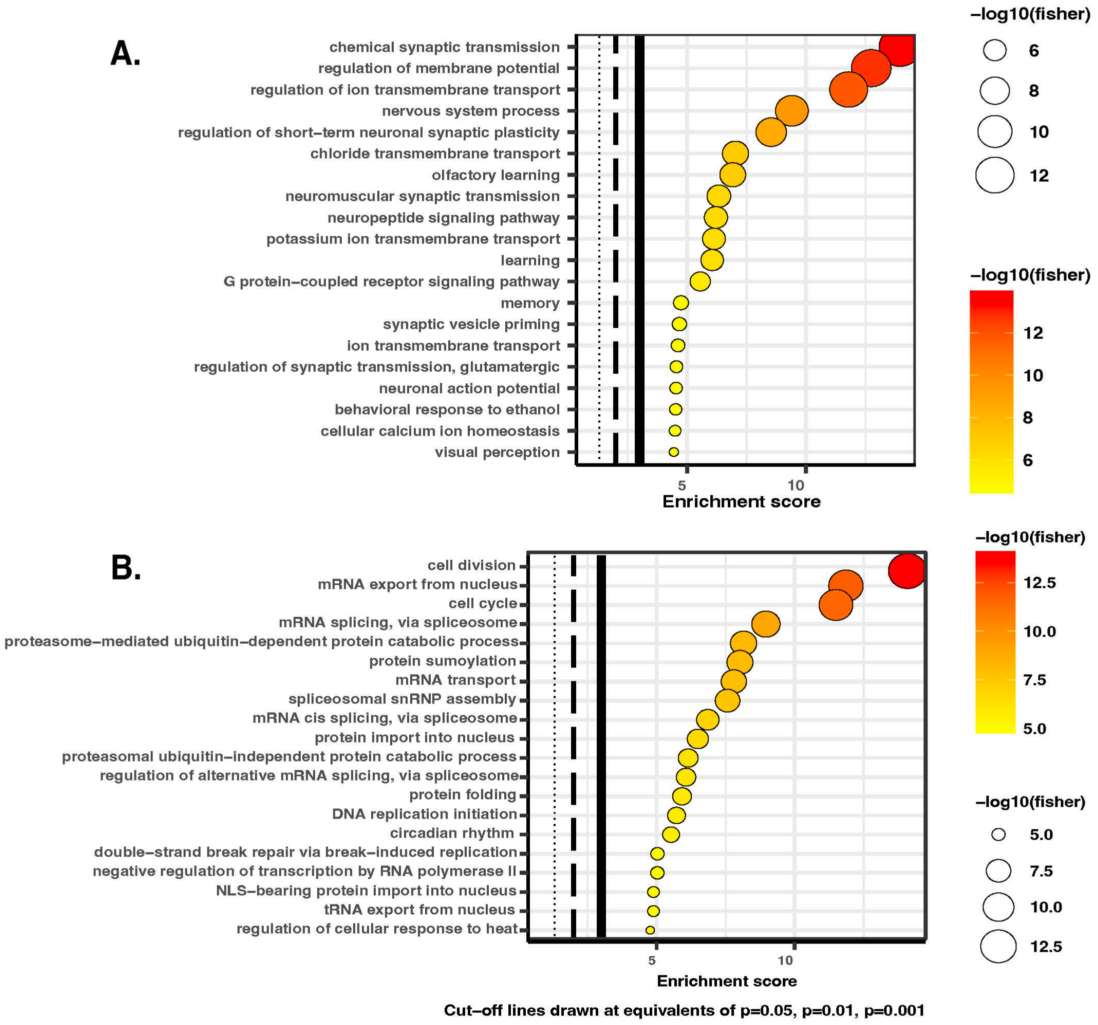


**Figure S14. Significant enrichment for GO Biological Processes, pupae brains.** A. GO enrichments for brain DEGs upregulated in social-biased vs solitary-biased pupae. B. GO enrichments for brain DEGs downregulated in solitary-biased pupae. Results are visualized using TopGO. Circle diameter and color is proportional to a -log10 pvalue from a Fisher Exact Test for enrichment of each GO term. Three significance threshold lines are included, for $p<0.05$ (dotted), $p<0.01$ (dashed), and $p<0.001$ (solid).

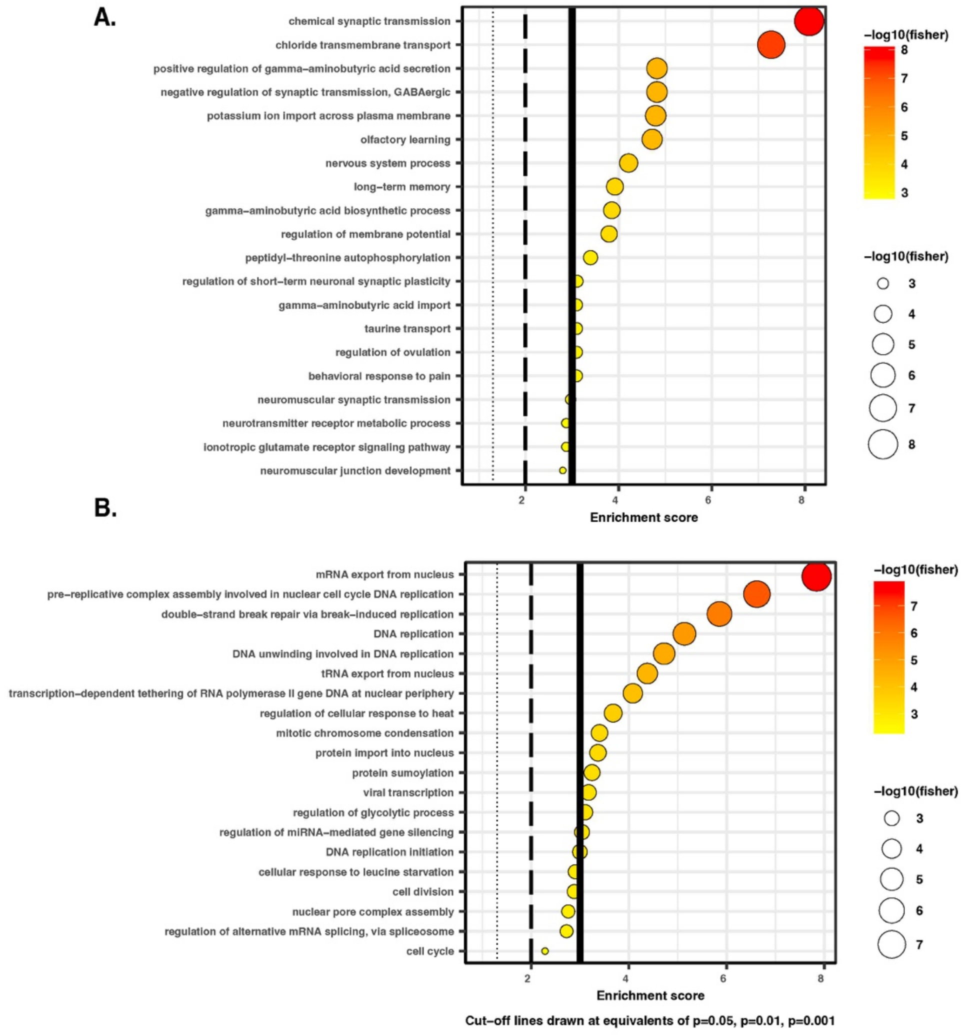


**Figure S15**. TopGO concordant DEGs, in pupal brains. A. GO enrichment for social-biased pupa genes. B. GO enrichment for solitary-biased pupa genes

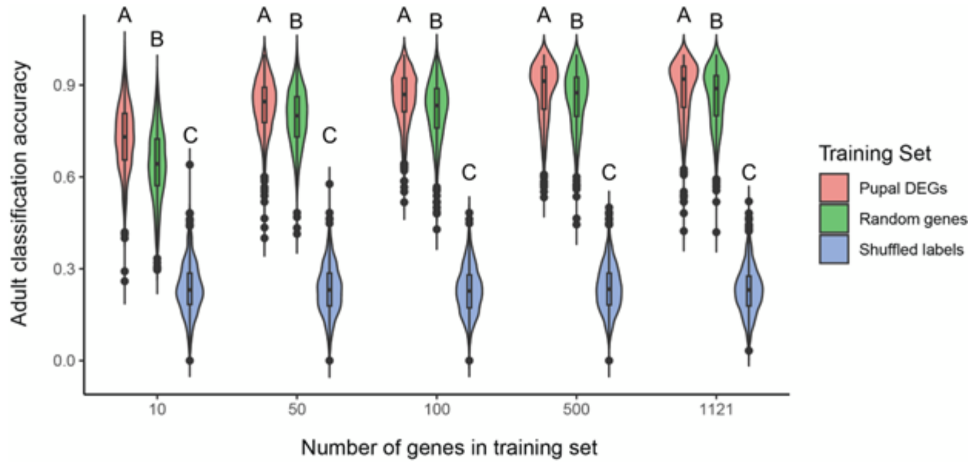


**Figure S16. Pupal brain differentially expressed genes accurately classify adult behavioral state.** A random forest model more accurately classified adult behavioral state using either subsets of or the entire list of 1121 pupal brain differentially expressed genes ("pupal DEGs") consistently performed better than a random, equally sized gene list that did not contain the pupal DEGs. A null control in which adult labels were shuffled ("Shuffled labels") consistently performed as well as chance alone (~25%), expectedly, as there are four adult behavioral groups. Violin plots show raw data as points with surrounding smoothed density curve; boxplots within show median as solid center line with first and third quartiles below and above, respectively. Letters denote significant differences (Scheffe's post-hoc test following ANOVA, $p<1e-12$ for each comparison).

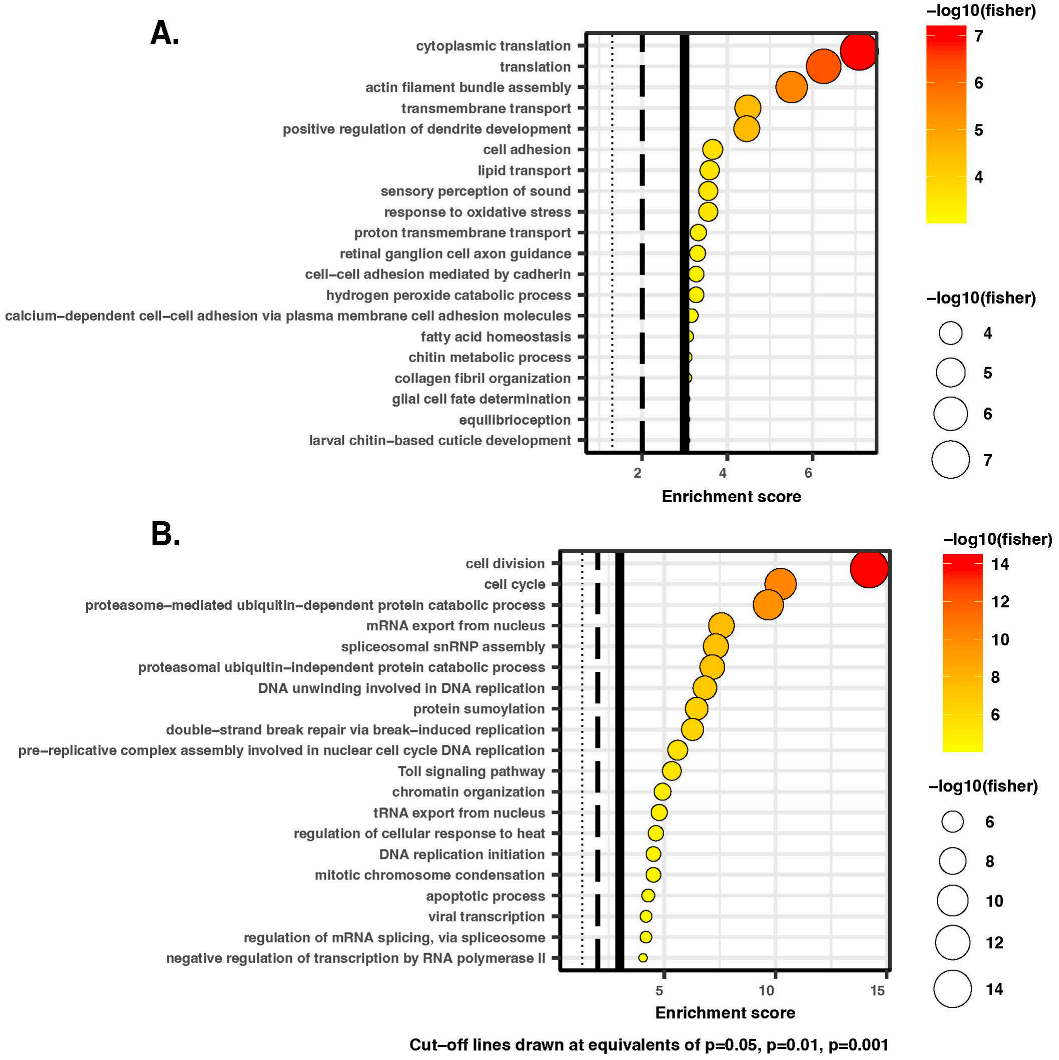


**Figure S17. Significant enrichment for GO Biological Processes, pupae fat bodies**. A. GO enrichments for fat body DEGs upregulated social-biased vs solitary-biased pupae. B. GO enrichments for fat body DEGs downregulated in solitary-biased pupae. Results are visualized using TopGO. Circle diameter and color is proportional to a -log10 pvalue from a Fisher Exact Test for enrichment of each GO term. Three significance threshold lines are included, for $p<0.05$ (dotted), $p<0.01$ (dashed), and $p<0.001$ (solid).

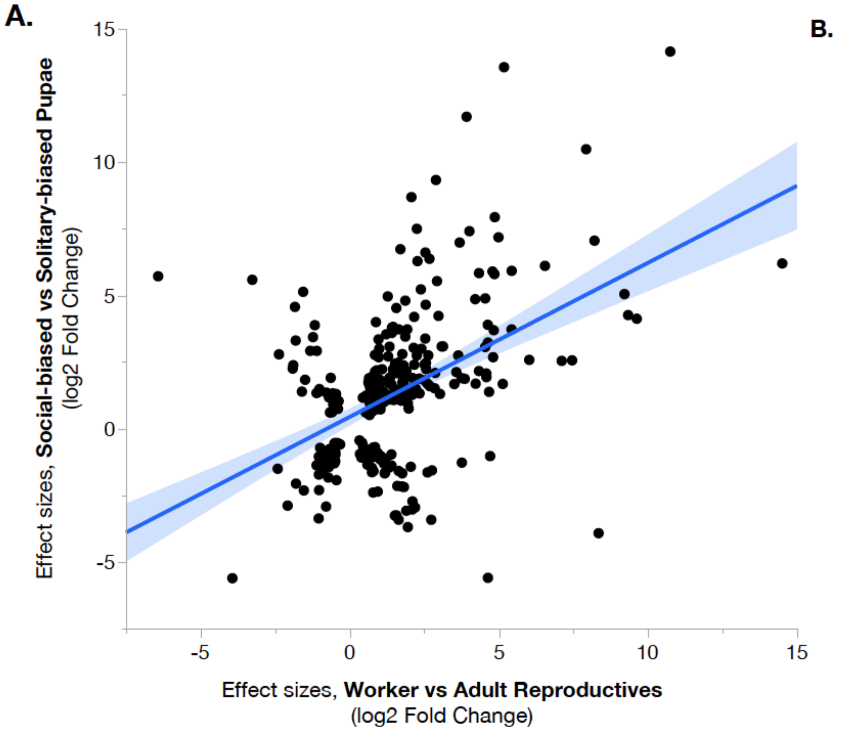


**B.**

| | Up, Social-Biased Pupae | Up, Solitary-Biased Pupae |
|---|---|---|
| Up, Adult Workers | 172***<br>Odds Ratio = 2.90 | 33 (ns)<br>Odds Ratio = 0.57 |
| Up, Adult Queens & Sol. Reprod. | 62 (ns)<br>Odds Ratio = 0.97 | 54 (ns)<br>Odds Ratio = 1.26 |

Odds Ratio 5 0

**Figure S18. Overlapping DEGs for adult and pupal fat body transcriptomes are correlated in direction and magnitude.** (A) The effect size estimates (shown here as the $\log_2$-fold change; DESeq2, FDR<0.001) for overlapping brain DEGs between adult reproductives vs. workers and solitary-biased vs. social-biased pupae are strongly correlated (linear regression, $R^2$=0.225, F=92.36, p=2.27e-19). (B) Genes upregulated in social-biased pupal fat bodies are enriched for genes that are also upregulated in adult workers (hypergeometric test, p=8.41e-26, odds ratio=2.90). Unlike in brains, genes more highly expressed in solitary-biased pupae are not more likely to also be upregulated in adult queens and solitary reproductives (hypergeometric test, p=0.07, odds ratio=1.26). There is no significant enrichment for discordant patterns of gene expression (e.g., genes upregulated in workers do not also show an enrichment for genes upregulated in solitary-biased pupae, etc). Asterisks denote a significant enrichment with p<0.0001.
